# Multiscale biological interactions define clinical trajectories in acute myeloid leukemia

**DOI:** 10.64898/2026.08.25.746926

**Authors:** Jan Bařinka, Sarah Gräßle, Magdalena Pajonk, Rosa Allesøe, Stefanos Bamopoulos, Maximilian Mönnig, Caroline Röthemeier, Lea Jopp-Saile, Dominik Vonficht, Eleni Besiridou, Jana Ihlow, Jan Braune, Evi Vlachou, Sergi Beneyto-Calabuig, Michael Kardorff, Selina Neunhäuser, Vincent Fregona, Simone Feurstein, Adriane Halik, Helen George, Judith Zaugg, Daniel Hübschmann, Michael Hundemer, Christoph Lutz, Andreas Trumpp, Lars Bullinger, Ulrich Keller, Carsten Müller-Tidow, Jörg Westermann, Frederik Damm, Jan Krönke, Tim Sauer, Lars Velten, Simon Raffel, Simon Haas

## Abstract

Cancer is characterized by complex interactions across genetic, cellular, and microenvironmental scales. However, a quantitative understanding of how these interactions shape clinical trajectories remains limited. Here, we present a multi-scale single-cell dataset from 184 treatment-naïve acute myeloid leukemia (AML) patients spanning all major genetic subtypes, together with an analytical framework to dissect interactions across biological scales. We show that distinct clinical outcomes are encoded by specific cross-scale, cross-compartment interactions present at diagnosis: response to induction therapy is governed by interactions between genetic alterations and leukemic differentiation state; relapse following chemotherapy is associated with non-genetic programs linked to metabolism; and relapse after allogeneic stem cell transplantation is driven by interactions between the immune microenvironment and residual healthy hematopoiesis. Together, our study provides a framework to resolve intra- and inter-patient heterogeneity in cancer and supports a model in which clinical trajectories in AML emerge from defined interactions across biological scales.

## INTRODUCTION

Cancer is increasingly recognized as a multi-layered disease in which clinical trajectories result from the interplay of genomic alterations, cancer cell states, and the surrounding micro- and macroenvironment^1^. Large-scale sequencing efforts have identified key driver mutations, patterns of clonal evolution, and genomic risk stratification frameworks^2,3^, while single-cell technologies have enabled high-resolution mapping of tumor cell states and microenvironmental interactions^4–6^. However, these advances have largely been developed in isolation. Cohort-level, single-cell multi-scale datasets that jointly capture these layers, together with quantitative frameworks to interrogate their interactions across molecular, cellular, tissue, and clinical scales, remain limited. As a result, a comprehensive understanding of how these layers interact to shape disease phenotypes and clinical trajectories remains a central challenge in cancer research.

Acute myeloid leukemia (AML) provides a paradigmatic cancer for investigating multi-layered interactions and their influence on disease trajectories^7–15^. Arising from transformed hematopoietic stem and progenitor cells (HSPC) in the bone marrow, AML exhibits pronounced inter- and intra-patient heterogeneity across genomic, cellular, and microenvironmental dimensions, reflected in highly variable clinical outcomes. To date, these layers have been studied largely in isolation. Large-scale genomic analyses have defined recurrent mutations and established widely used classification systems^11,12,16–19^, while longitudinal bulk and single-cell multi-omic studies have characterized clonal evolution and molecular programs associated with progression and treatment response^11,13,16,20–27^. In parallel, non-genetic features, including leukemic stem cell differentiation states, have emerged as key determinants of disease behavior^28–32^. At the same time, the bone marrow microenvironment, including the immune system, has been shown to critically influence disease progression and therapeutic response^14,33–35^. As a result, existing classification systems – whether genomic, stem cell–centric, or microenvironmental – remain largely disjointed, capturing only partial aspects of disease biology and failing to fully explain clinical trajectories at diagnosis. Despite emerging efforts to connect individual layers of AML heterogeneity^13,20–22,28,32,33^, a comprehensive and integrated understanding of how diverse biological scales interact to shape disease phenotypes and clinical trajectories remains lacking. Here, we address this gap by presenting a cohort-level, multi-scale single-cell dataset of 184 treatment-naïve AML patients, together with a quantitative analytical framework to interrogate interactions across biological scales, cellular compartments, and clinical outcomes. We define AML archetypes based on recurrent genotype–phenotype relationships in leukemic stem and progenitor cells, identify genotype-driven traits alongside non-genetic metabolic programs as an independent axis of heterogeneity, and delineate how these leukemia-intrinsic features interact with the immune microenvironment and residual healthy hematopoiesis. Our findings demonstrate that distinct clinical outcomes arise from specific combinations of mechanisms operating across biological scales that are already present at diagnosis, providing a unified framework to understand the biological basis and clinical consequences of AML heterogeneity.

## RESULTS

### A multi-scale single-cell framework of acute myeloid leukemia

To establish robust associations between genomic aberrations, leukemic cell states, transcriptomic programs, surface markers, the microenvironment, and clinical outcomes, we generated a multi-scale single-cell dataset from 184 therapy naïve AML patients and eight healthy controls, encompassing all major subtypes according to WHO 2022^17^ and ICC 2022^18^, as well as recurrent genetic driver aberrations (Figure 1a,b; Supplementary Figure 1, Supplementary Table 1). This dataset comprises: (i) single-cell proteo-genomic data combining transcriptomic sequencing and 81 surface proteins (n=184 patients; 187,418 cells); (ii) ultra-high-parametric cytometry-based phenotyping of the cellular composition of the AML immune microenvironment (n=152; 41,5 million cells); (iii) targeted genotyping of recurrent AML driver mutations (n=184); and (iv) detailed clinical outcome data alongside standard diagnostic information (n=184). The investigated subtypes include: AML, myelodysplasia-related (n=62), AML with NPM1 mutation (n=45), with RUNX1::RUNX1T1 fusion (n=14), with CEBPA mutation (n=10), with CBFB::MYH11 fusion (n=10), with MECOM rearrangement (n=9), with PML::RARA fusion (n=9), with KMT2A rearrangement (n=5), with DEK::NUP214 fusion (n=2), not otherwise specified, defined by differentiation (n=18) and healthy donors (n =8) (Figure 1b, Supplementary Figure 1). Importantly, the observed clinical course across the investigated cohort and the AML subtypes closely mirrored prior studies, supporting the cohort’s representative character (Figure 1b, Supplementary Figure 2a).

**Figure 1.**
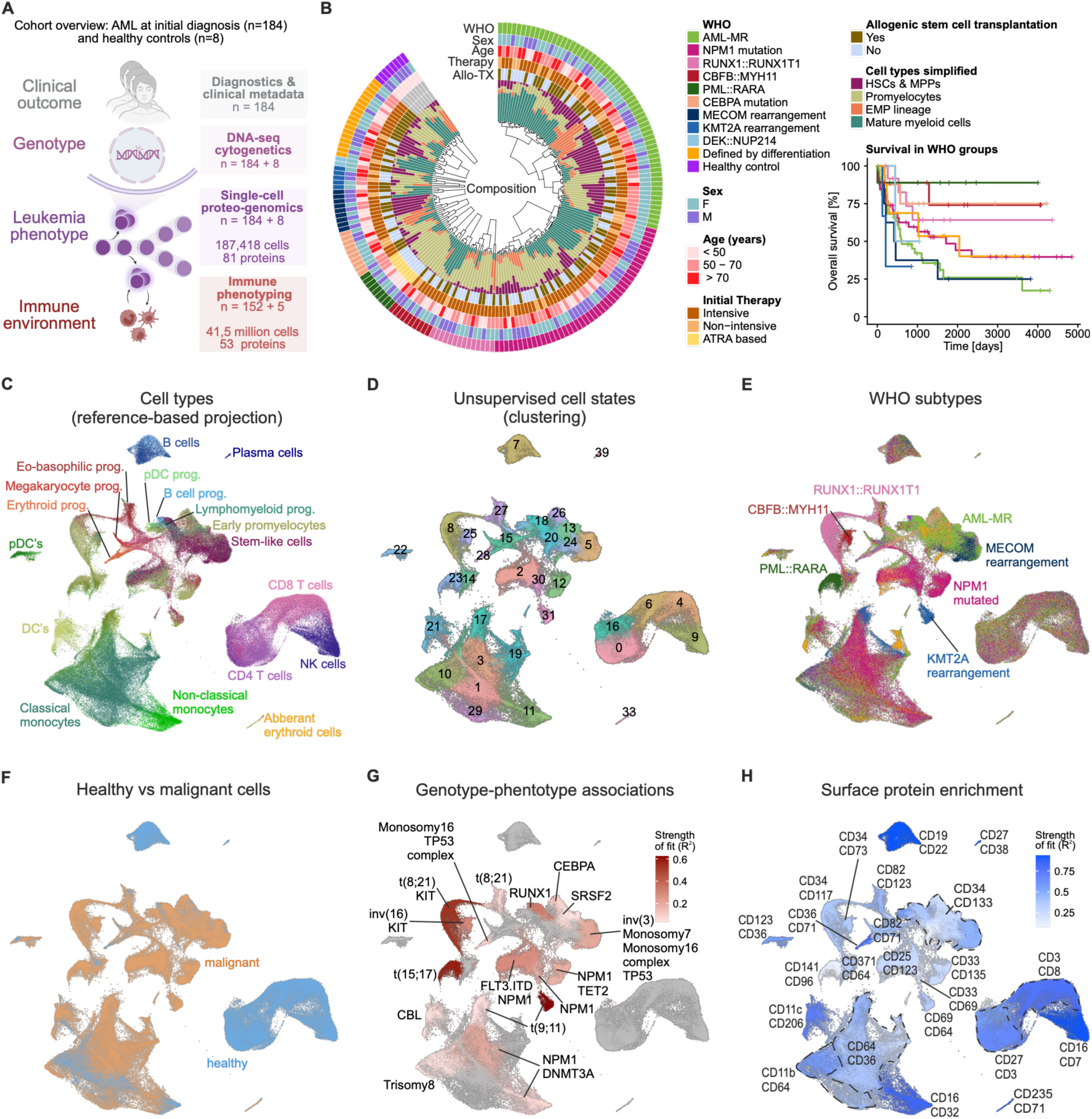
The cellular landscape of therapy-naïve AML across genetic subtypes. **(A)** Schematic overview of the cohort and acquired data modalities. **(B)** Selected patient metadata and simplified compositional overview. The inner circle depicts the distribution of major myeloid cell types derived from single-cell transcriptomic data. Patients were hierarchically clustered within WHO-defined groups. Bottom right: overall survival of the cohort stratified by WHO subtypes. A comprehensive overview of metadata is provided in Supplementary Figure 1. **(C–H)** UMAP embeddings of single-cell transcriptomic data. **(C)** UMAP colored by projected cell types and differentiation states based on label transfer from a healthy reference^35^. **(D)** UMAP colored by cell states identified via unsupervised Louvain clustering. **(E)** UMAP colored by WHO 2022 subtypes. **(F)** UMAP colored by healthy versus malignant status (see methods). **(G)** UMAP colored by genotype–phenotype associations, showing enrichment of patient-level genomic aberrations across Louvain clusters. Mutation enrichment per cluster was assessed using univariate negative binomial generalized linear models. Only aberrations with q < 0.01 and positive effect size (β > 0) are shown. **(H)** UMAP colored by enrichment of surface protein expression (AbSeq). Marker expression was modeled using linear mixed-effects models across Louvain clusters. Cluster- specific contrasts versus the mean of all other clusters were computed, and the top two markers per cluster (by effect size) are displayed. **(G–H)** The overall explanatory power of selected mutations **(G)** and surface markers **(H)** per cluster is reported as adjusted McFadden’s pseudo- R², with higher values indicating stronger genotype–transcriptome **(G)** or proteome– transcriptome **(H)** associations.

UMAP visualization of single-cell transcriptomic data, followed by projection onto a healthy hematopoietic reference^35^, enabled cell type annotation and positioned both normal and leukemic stem and progenitor cells along differentiation trajectories (Figure 1c, Supplementary Figure 2b). Notably, UMAP embedding of the surface proteomic layer resolved cell type at a highly comparable level, indicating that the surface marker space effectively captures the underlying cellular identity (Supplementary Figure 2c). Unsupervised clustering of the single-cell transcriptomic data defined a comprehensive map of leukemic and microenvironmental cell states in treatment-naïve AML (Figure 1d). Importantly, these unsupervised transcriptomic cell states outperformed projected cell type and differentiation state information in explaining WHO subtypes, highlighting the additional biological and clinical value of our approach over previous bulk deconvolution and single-cell reference-based approaches^28,29^ (Figure 1e; Supplementary Figure 2d,e).

To distinguish healthy from leukemic cells, we determined NPM1 mutation status and copy number variations (CNVs) directly from single-cell transcriptomic data (Supplementary Figure 3a– e). Using this information as ground truth, we developed and validated a model that predicts malignancy status in the HSPC compartment with high accuracy based on transcriptional features alone (Supplementary Figure 3f, see methods for details). This model was subsequently applied to all cells lacking genetic ground truth (Figure 1f). Indeed, the proportion of inferred leukemic cells correlated with blast counts from routine bone marrow diagnostics (Supplementary Figure 3g), and residual healthy cells from AML patients co-clustered with cells from healthy individuals along the normal stem cell differentiation trajectory and within B, T, and NK cell compartments (Figure 1f). In contrast, leukemic cells from patients converged toward distinct transcriptomic states in the stem and progenitor compartment, consistent with strong genotype– phenotype relationships. To systematically delineate these relationships, we quantified associations between genomic aberrations and recurrent leukemic cell states (Figure 1g; methods). While some established AML drivers, such as the translocations t(15;17) (PML::RARA), t(8;21) (RUNX1::RUNX1T1), and t(9;11) (KMT2A::MLLT3), showed strong associations with defined transcriptomic states in the stem and progenitor compartment, others, including NPM1, DNMT3A and TET2 mutations, exhibited weaker associations (Figure 1g). Using a similar strategy, we further mapped associations between leukemic cell states and their surface proteomic profiles (Figure 1h, Supplementary Table 2). A list of surface markers associated with cell type identities and WHO subtypes is provided in the Supplementary Table 3.

To quantitatively characterize the cellular composition of the AML immune microenvironment, we employed three high-dimensional cytometry panels^36^ designed to comprehensively profile immune cell diversity and states (Supplementary Figure 4a–c, Supplementary Table 2). These included: (i) a broad immune profiling panel to identify major immune cell populations, including B cell, T cell, and NK cell subsets; (ii) a T cell focused panel to resolve the phenotypic states of CD4, CD8 αβ, and γδ T cells; and (iii) a myeloid-focused panel to finely map AML differentiation trajectories. Integrative analysis of these panels enabled the quantification of 34 distinct immune cell types and states across more than 41 million cells (Supplementary Figure 4d,e, Supplementary Table 1).

To systematically quantify disease characteristics across biological scales, we computed a set of interpretable features for each patient (see methods). These include (i) the differentiation block of leukemic cells, defined as the mean pseudotime of stem and progenitor differentiation; (ii) the fraction of leukemic cells exhibiting transcriptomic features of hematopoietic stem cells (phenotypic leukemic stem cells; % pLSCs); (iii) the average cell cycle activity within the phenotypic leukemic stem and progenitor cell (pLSPC); compartment (iv) the degree of residual healthy hematopoiesis, as defined by proportion of healthy versus malignant cells within stem and progenitor compartment; (v) the transcriptomic similarity of leukemic cells to healthy cells (healthy-like score; see methods, Supplementary Figure 5a); (vi) the cellular composition of the immune microenvironment; and (vii) the activity of gene expression programs across cellular compartments, as inferred from outcome-based gene set enrichment analyses (GSEA), alongside commonly used transcriptomic scores, such as LSC17 score^31^. All derived features are fully accessible in Supplementary Table 1, alongside patient-level metadata including age, WHO classification, ELN risk, and clinical outcome.

Together, this dataset provides a unified view of leukemic and immune microenvironmental states across genetic subtypes, enabling systematic interrogation of interactions across cellular compartments and biological scales.

### Combinatorial architecture of biological variation underlying clinical trajectories in AML

To dissect how distinct biological layers at diagnosis jointly shape clinical trajectories, we developed an analytical framework that first quantifies relationships between sources of inter-patient variation (Figure 2a) and then evaluates their associations with clinical outcomes (Figure 2b).

**Figure 2.**
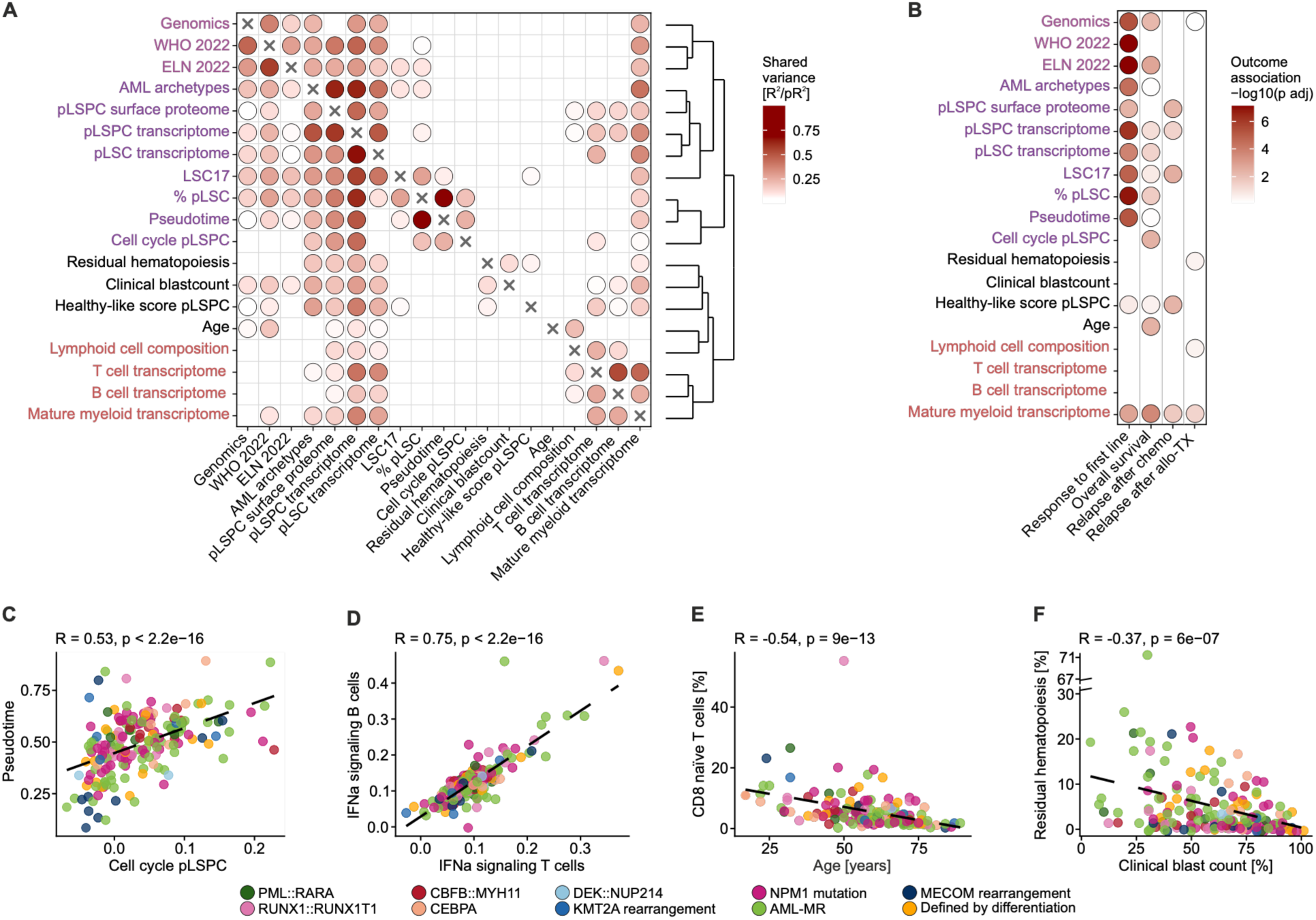
Combinatorial Structure of Biological Variation Underlying Clinical Trajectories in AML. **(A)** Dot plot summarizing pairwise predictive relationships between variables and variable groups. Rows represent outcomes and columns represent predictors, modeled using generalized linear frameworks. Continuous outcomes were modeled by linear regression (adjusted R²), categorical outcomes by logistic or multinomial regression (adjusted McFadden’s pseudo-R²), and multivariate numeric groups (e.g. transcriptomic or proteomic signatures represented by principal components) by multivariate linear models (multivariate R² trace-based). Statistical significance was assessed using likelihood ratio or F-tests. Dot color represents model explanatory strength (R² or pseudo-R², depending on outcome type). Only associations with adjusted p < 0.1 and adjusted R² > 0.05 are shown. Variables were hierarchically clustered based on reciprocal predictive strength. **(B)** Dot plot summarizing associations between variable groups in rows and clinical endpoints in columns. Response to induction chemotherapy was modeled using logistic regression, whereas overall survival and relapse endpoints were modeled using Cox proportional hazards regression. For time-to-event outcomes, proportional hazards violations were assessed and, where supported, time-varying effects were incorporated using penalized spline terms. Model significance was evaluated by likelihood ratio tests against null models. Dot color indicates significance as −log10 adjusted p-value. Only associations with adjusted p < 0.1 and improved model fit relative to the null model (ΔAIC > 0) are shown. **(A,B)** P-values were adjusted using the Benjamini–Hochberg method. **(C–F)** Selected pairwise associations highlighting key relationships identified in Fig. 2A. Spearman’s rank correlation coefficient is indicated. Each point represents an individual patient sample, colored by WHO 2022 classification. Dashed lines show ordinary least squares fits for visualization. **(C)** Relationship between mean cell cycle activity and the differentiation state (pseudotime) in phenotypic leukemic stem and progenitor cells (pLSPCs). **(D)** Association of interferon-α (IFNα) gene set expression in T and B cells within the AML microenvironment. **(E)** Association between the proportion of CD8 naïve T cells and patient age. **(F)** Relationship between clinical blast counts and the extent of residual healthy hematopoiesis.

We first quantified how much variation in one biological layer (e.g. genetics) can explain variation in another (e.g. cell states, microenvironment etc.) using a coefficient of determination (R²)- based metrics, thereby identifying shared versus independent sources of heterogeneity (Figure 2a; see methods). High values indicate that variation in one feature group can be largely explained by another, suggesting overlapping information content, whereas low values indicate that feature groups capture distinct aspects of patient variability. Clustering in the shared information space identified two major axes representing leukemia-intrinsic and leukemia- extrinsic variation, which further subdivide into smaller sub-branches of features with overlapping variance (Figure 2a).

Within the leukemia-intrinsic axis, three sub-branches emerged: (i) genetic features (genomics, WHO and ELN22 subtypes), (ii) transcriptomic and proteomic features of leukemic populations, and (iii) features related to leukemic cell cycle activity and differentiation state (Figure 2a). While genetic features showed minimal shared information with microenvironmental features, transcriptomic and surface proteomic profiles of leukemic populations shared information across both axes, consistent with them receiving signals from both cell-intrinsic and extrinsic sources (Figure 2a). Notably, cell cycle activity in pLSPCs was strongly associated with differentiation state, as captured by pseudotime and the proportion of immature pLSCs (Figure 2a,c). This relationship was partly linked to genetic subtype, consistent with a preferential genotype-dependent arrest of malignant HSPCs at distinct differentiation states across subtypes (Supplementary Figure 5b–d). Despite these genotype–phenotype associations, a large fraction of leukemia-intrinsic transcriptomic variation, including key biological processes such as metabolism and inflammation, is not explained by genotype (Supplementary Figure 5e), as we detail below.

The second principal axis comprised immune microenvironmental features on one side and features related to disease burden on the other (Figure 2a). Within the microenvironmental sub-branch, distinct immune cell types, such as T cells, B cells, and mature myeloid cells, encoded shared transcriptomic information. For example, type I interferon responses were highly correlated across immune cell types, likely reflecting shared exposure to systemic cues in the AML microenvironment (Figure 2d). The composition of the non-malignant lymphoid compartment was closely linked to these programs but also associated with patient age (Figure 2a), consistent with age-related immune remodeling, such as a decline of naïve T cells (Figure 2e). In the other sub-branch, disease burden–related features clustered together, with blast counts from routine diagnostics being inversely correlated with both residual hematopoiesis and the healthy-like score (Figure 2f; Supplementary Figure 5f).

Notably, distinct clinical outcomes were associated with different axes and sub-branches (Figure 2b). Response to induction chemotherapy was primarily associated with leukemia- intrinsic features, including genomic subtype and pLSPC differentiation state. Relapse after chemotherapy was linked to non-genetic leukemia-intrinsic heterogeneity, whereas relapse following allogeneic stem cell transplantation was associated with leukemia-extrinsic factors, such as the immune system and the extent of residual healthy hematopoiesis – all features not captured by current classification systems or risk scores. As expected, overall survival was determined by a combination of these features, with additional overarching factors, such as leukemic cell cycle and patient age, also contributing.

Collectively, this framework delineates intra- and inter-patient heterogeneity and reveals how distinct leukemia-intrinsic and microenvironmental factors present at diagnosis jointly shape clinical outcomes, as further detailed below.

### AML archetypes capture recurrent leukemic stem and progenitor cell–intrinsic genotype– phenotype relationships

We first focused on leukemia-intrinsic variation. Although genomic aberrations and their complex combinations have been extensively characterized in AML, their relationship to leukemia phenotypes has primarily been studied using bulk gene expression profiles, obscuring differential effects of genotype on distinct cell types and differentiation stages. To address this limitation, we leveraged the single-cell nature of our data to examine in which cellular compartments genotype–phenotype associations are most pronounced (Figure 3a). The strongest association between genomic alterations and recurrent transcriptomic phenotypes was observed within the broader pLSPC compartment, defined by stem- and progenitor-like differentiation states (Figure 3a). This association was markedly attenuated both in mature myeloid leukemic cells downstream of the differentiation block and when analyses were restricted to phenotypic leukemic stem cells (pLSCs) (Figure 3a). This suggests that genotype–phenotype relationships across AML subtypes are most robustly captured in the broader stem and progenitor compartment, rather than in fully differentiated or strictly stem-like subsets.

**Figure 3.**
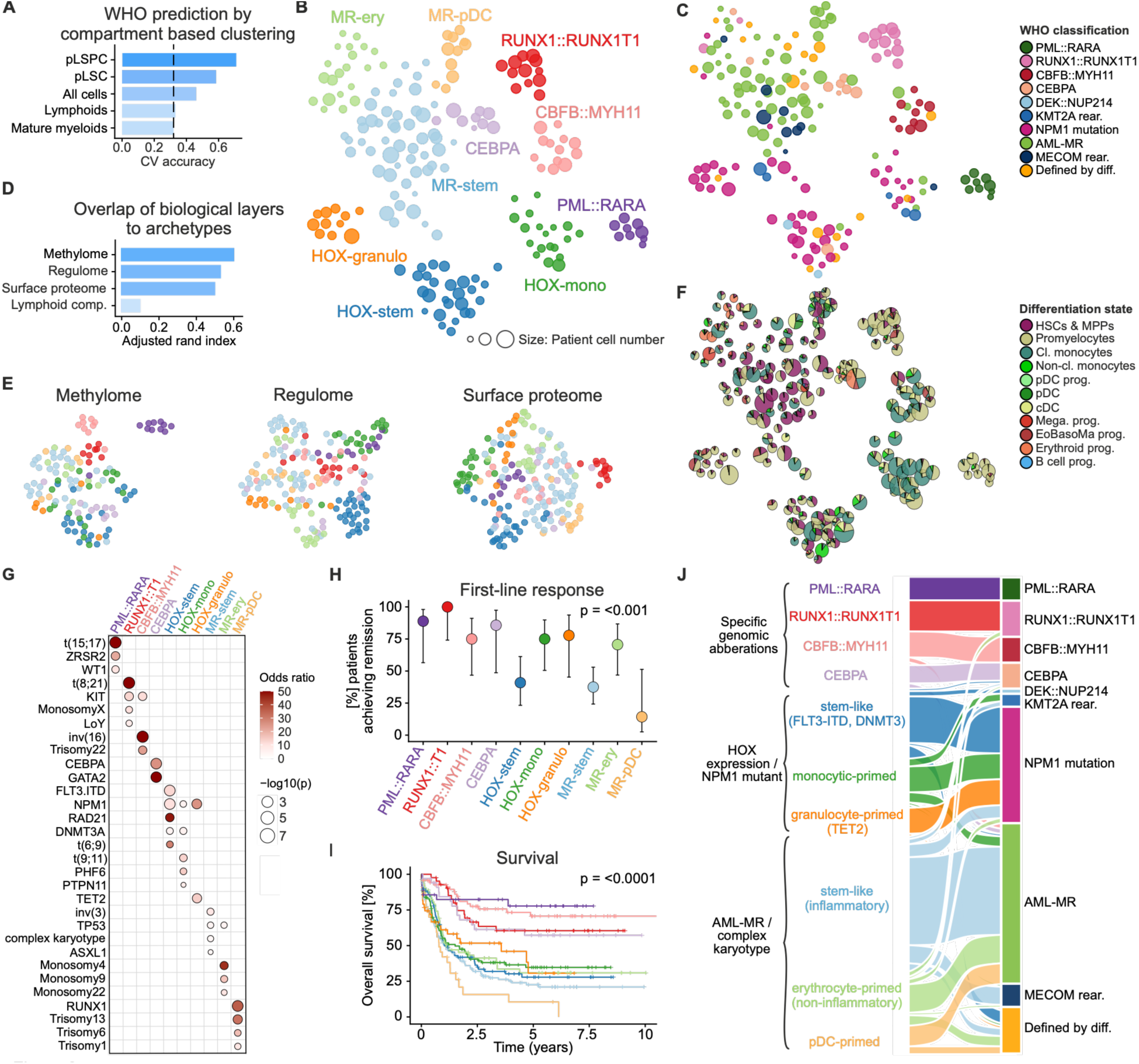
Leukemic stem and progenitor cell–based classification defines AML archetypes. **(A)** Association between cellular compartments and genetic WHO 2022 subtypes. To assess how well distinct cellular compartments can explain WHO 2022 subtypes, for each compartment patient- level pseudobulk were generated and clustered into groups. WHO 2022 classification was then predicted using multinomial logistic regression, and performance was evaluated by 5-fold cross- validation. Bars indicate mean cross-validated classification accuracy. **(B)** UMAP colored by AML archetypes. Archetypes and UMAP embedding are based on pLSPC transcriptome pseudobulks, using 1000 most variable genes. **(C)** AML archetype UMAP colored by WHO 2022 classification. **(D)** Relationship between transcriptome-based AML archetypes and other molecular layers quantified by Rand index. Regulome layer is based on the results of SCENIC^38^ within the pLSPC compartment, surface proteome is based on surface proteins quantified by Abseq, methylome clustering overlap was evaluated on methylation array data available for the TCGA dataset^12^, where AML archetype labels were predicted prior to determining the overlap (methods). The lymphoid layer was defined by lymphoid cell composition based on flow cytometry data. **(E)** UMAP embeddings based on the patient-level datasets for the respective data layers and colored by AML archetypes. **(F)** AML archetype UMAP showing cell type composition, derived from the single-cell transcriptome data. Pie charts represent cell type composition within each sample. **(G)** Enrichment of genetic aberrations across AML archetypes. For each genetic aberration and archetype, enrichment was assessed by comparing the frequency in a given archetype versus all other archetypes combined using odds ratios from 2 × 2 contingency tables. P-values were calculated from the same contingency tables and adjusted using the Benjamini–Hochberg method. Only enriched associations with odds ratio > 1 and adjusted p < 0.1 are shown. **(H)** Response to induction therapy across AML archetypes. Percentage of patients achieving complete remission is indicated, with error bars indicating 95 % confidence intervals calculated using the Wilson method. Overall differences across archetypes were assessed using Fisher’s exact test based on contingency tables of response categories. **(I)** Overall survival (OS) in AML archetypes. To increase patient numbers, AML archetype identities were determined in publicly available bulk-RNAseq dataset (n=727)^37^ with available OS data (methods). **(J)** Alluvial plot showing relationship between AML archetypes (left) and WHO 2022 classification (right). **(B,C,F)** Dot size is scaled by cell number per patient in pLSPC compartment.

To derive a classification that optimally captures these recurrent genotype–phenotype states, we performed clustering analyses on patient-level gene expression summaries (pseudobulks) of leukemic stem and progenitor cells, explicitly excluding both microenvironmental and mature leukemia-derived cells. This approach identified ten distinct clusters, in the following referred to as AML archetypes (Figure 3b). These archetypes were associated with specific genotypes (Figure 3c,g), DNA methylation landscapes (inferred from bulk data^37^; methods), gene regulatory programs (SCENIC based^38^), surface proteomic profiles (Figure 3d,e), and differentiation states (Figure 3f). Importantly, AML archetypes showed a highly significant association with response to induction therapy and overall survival (Figure 3h,i). AML archetypes were largely independent of microenvironmental immune cell composition (Figure 3d), indicating that these clusters represent leukemia-intrinsic core states that integrate recurrent genetic, epigenetic, transcriptional, and phenotypic inter-patient variation to defined subtypes that are linked to clinical outcome.

Importantly, many AML archetypes closely corresponded to established WHO-defined genomic AML subtypes (Figure 3c,j). In particular, subtypes driven by single, dominant genomic events generally exhibited the most pronounced genotype-dependent transcriptional, epigenomic, and surface proteomic programs within leukemic stem and progenitor cells and formed homogeneous clusters consistent with current genomic classifications (Figure 3b,g). These included CBFB::MYH11 AML, RUNX1::RUNX1T1 AML, PML::RARA AML, and AML with CEBPA mutations. In contrast, several other WHO subtypes displayed substantial inter-patient heterogeneity within the leukemic stem and progenitor compartment that is not fully captured by existing genomic classification schemes. For example, NPM1-mutated AMLs alongside other AMLs segregated into three distinct clusters (Figure 3b,c,j). These were characterized by high HOX gene expression and designated based on their differentiation states and enriched co-mutation patterns (Supplementary Figure 6a; Figure 3g,j). Similarly, myelodysplasia-related (MR) AML cases partitioned into three major groups defined by recurrent genomic alterations, differentiation states and biological features, which will be detailed in the following section.

Collectively, these findings provide a comprehensive framework for understanding how genomic alterations converge on stable leukemic stem and progenitor cell states and establish a refined AML classification based on recurrent genotype–phenotype relationships.

### Cellular architecture and biological traits of AML archetypes

To define the cellular architecture of AML archetypes, we characterized their differentiation blocks, assessed transcriptomic lineage priming toward all hematopoietic lineages, and derived archetype-specific surface marker profiles (Figure 4a, Supplementary Table 3). To identify biological processes associated with each archetype, we performed GSEA while accounting for differences attributable to distinct differentiation states (Figure 4b, see methods). These analyses revealed that each archetype exhibited a unique differentiation block, with distinct lineage priming toward monocytic, neutrophilic, eosinophil–basophil, plasmacytoid dendritic cell (pDC), erythroid, megakaryocytic or early B cell lineages, translating into archetype-specific traits.

**Figure 4.**
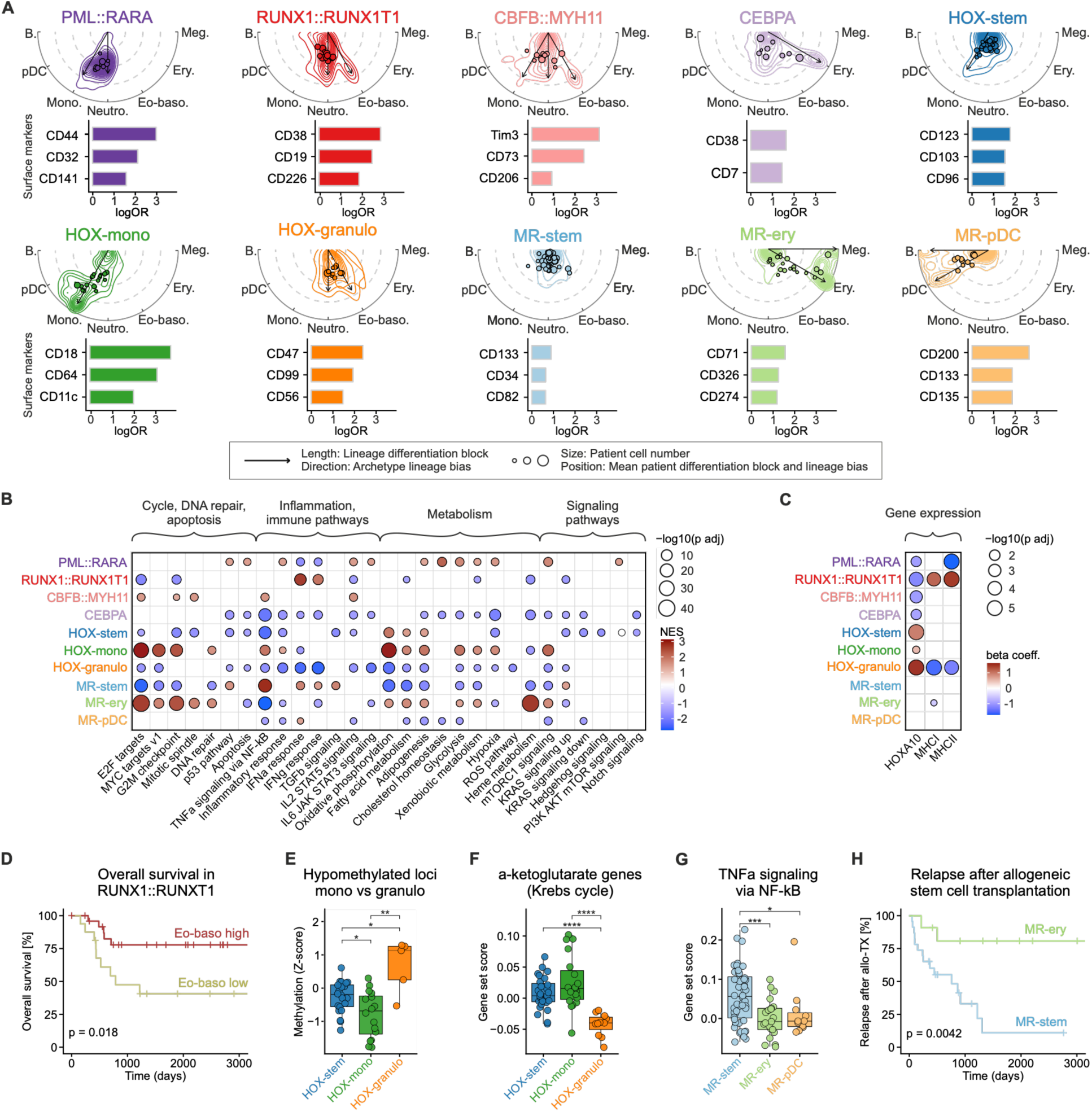
Cellular architecture and biological traits of leukemic stem cell archetypes. **(A)** Cellular differentiation (top rows) and surface marker associations (bottom rows) of AML archetypes. Top rows: Semi-circular plots summarize differentiation trajectories in AML archetypes. Radial position reflects differentiation (pseudotime), and angular position reflects lineage bias. Dots indicate patient-level average positions, with dot size proportional to the number of cells contributing to each patient average. Contours represent patient-balanced two-dimensional density estimates averaged within each archetype. Arrows denote the major lineage directions for each archetype, with arrow length reflecting the extent of differentiation along the respective lineage. Bottom rows: Surface marker associations for AML archetypes. For each marker– archetype combination, association was assessed using Firth’s bias-reduced logistic regression. Marker expression was averaged per-patient in the pLSPC compartment and up to three most significant positively associated markers per archetype are shown. Bar length represents the standardized log odds ratio (logOR). **(B)** Hallmark pathway enrichment across AML archetypes. pLSPC pseudobulks were analyzed by DESeq2 in a one-versus-rest design with cell type compositions as covariates. Genes were ranked for each archetype and analyzed by geneset enrichment analyses (GSEA) of MSigDB hallmark gene sets. Hallmarks without clear biological interpretability in this context, such as spermatogenesis, estrogen response, or UV response pathways, were excluded from visualization. Only pathways with adjusted p < 0.05 are shown. Dot color represents normalized enrichment scores (NES), and dot size represents −log10 adjusted p-value. **(C)** Gene expression differences across AML archetypes. For each gene, standardized expression was modeled using linear regression with archetype membership (one- versus-rest) and progenitor cell type composition as covariates. Dot color represents the regression coefficient for archetype membership, and dot size represents −log10 adjusted p- value. Associations with adjusted p < 0.1 are shown. **(D)** Overall survival in RUNX1::RUNX1T1 AML stratified by eosinophilic–basophilic cell proportion. To increase sample numbers, a publicly available bulk RNA-seq dataset^37^ was deconvoluted using dataset-derived signatures and the proportion of eosinophilic cells was determined. Patients were stratified by an outcome- optimized cutoff. **(E)** Monocyte–granulocyte lineage methylation across HOX archetypes. Differentially methylated CpG loci between healthy monocytes and granulocytes were identified from public reference methylation data^12^. A granulocyte-biased methylation score was computed in TCGA AML samples using the mean z-scored M-value across granulocyte-enriched loci and compared across predicted HOX archetypes. **(F)** Expression of alpha-ketoglutarate genes across HOX subtypes. Genes related to alpha-ketoglutarate generation in the Krebs cycle were aggregated into a geneset and quantified in pLSPC compartment across HOX archetypes. **(G)** Expression of the tumor necrosis factor alpha (TNFα) signaling pathway via nuclear factor kappa B (NF-κB) hallmark gene set in pLSPC compartment across myelodysplasia-related (MR) archetypes. **(H)** Relapse after allogeneic stem cell transplantation in MR-stem and MR-ery AML archetypes. **(A–C)** P-values were adjusted using the Benjamini–Hochberg method. **(E–G)** Group differences were assessed by Wilcoxon rank-sum tests. **(D, H)** Survival differences were assessed by log-rank tests. Significance levels are indicated as follows: * p < 0.05, ** p < 0.01, *** p < 0.001, **** p < 0.0001.

CBFB::MYH11 and RUNX1::RUNX1T1 leukemic stem and progenitor cell archetypes occupied distinct yet closely related transcriptional states, consistent with both rearrangements targeting subunits of the core-binding factor (CBF) transcriptional complex, a central regulator of normal hematopoiesis^39^ (Figure 4a, see also Figure 1e). In both subtypes, leukemic stem and progenitor cells formed a continuum extending from stem-like states toward early neutrophilic (promyelocytic) differentiation along one axis, and toward eosinophil–basophil differentiation along the other (Figure 4a, arrows; Figure 1e; Supplementary Figure 6b). In line with previous reports^40^, RUNX1::RUNX1T1 AMLs exhibited elevated tonic interferon signaling (Figure 4b), resulting in increased baseline expression of antigen presentation machinery, including MHC class I and class II molecules (Figure 4c). Within RUNX1::RUNX1T1 AML, eosinophil–basophil differentiation was marked by CD226 surface expression (Supplementary Figure 6b–d) and was associated with favorable overall survival in this subtype (Figure 4d). In contrast, leukemic progenitor cells in PML::RARA acute promyelocytic leukemia displayed a homogeneous differentiation block at the promyelocytic/neutrophilic stage (Figure 4a), with no detectable phenotypic stem cell compartment. Consistent with this neutrophilic arrest, these leukemias exhibited a distinct metabolic state, extensive hypermethylation around the transcription start site of CIITA – the master transcriptional regulator of MHC class II – and correspondingly low MHC class II expression, in line with HLA-DR negativity of this subtype in clinical routine diagnostics (Figure 4c, Supplementary Figure 6e).

The three AML archetypes defined by high HOX gene expression exhibited distinct differentiation states (Figure 4a, 3f), co-mutation patterns (Figure 3g), and biological characteristics (Figure 4b,c) and were designated Hox-stem, Hox-mono, and Hox-granulo according to their respective differentiation blocks. Hox-stem AMLs were enriched for NPM1 mutations co-occurring with mutations in DNMT3A, cohesin complex genes, and FLT3 internal tandem duplications (ITD), as well as cases harboring DEK::NUP214 fusions (Figure 3g), and were characterized by an arrest in a stem-like state with limited evidence of lineage commitment (Figure 4a). Consistent with its pronounced stemness features, this subtype demonstrated inferior responses to frontline chemotherapy (Figure 3h). Hox-granulo AMLs were enriched for NPM1 mutations in combination with TET2 or IDH1/2 mutations and exhibited a differentiation block at an early neutrophil- or eosinophil/basophil-primed progenitor stage (Figure 4a). This subtype displayed global DNA hypermethylation in line with the presence of TET2 and IDH1/2 mutations, well-established drivers of epigenetic hypermethylation in AML^41^ (Supplementary Figure 6f). Differentially methylated CpG regions associated with monocyte versus granulocyte differentiation were particularly affected, suggesting an epigenetic mechanism underlying the observed granulocytic lineage priming in this subtype (Figure 4e). For example, compared to other subtypes, Hox- granulo AMLs exhibited promoter hypermethylation of CIITA, thereby phenocopying the other granulocytic-primed subtype, PML::RARA AML (Supplementary Figure 6e). This was associated with reduced MHC-II expression, consistent with the lower MHC-II levels characteristic of granulocytic relative to monocytic cell states (Figure 4c). Notably, in addition to mutations affecting regulators of α-ketoglutarate metabolism (IDH1/2) or α ketoglutarate–dependent dioxygenases (TET2), genes involved in α-ketoglutarate biosynthesis were consistently downregulated, pointing toward a convergent metabolic dysregulation in this subtype (Figure 4f, Supplementary Figure 6g). Finally, Hox-mono AMLs comprised predominantly NPM1-mutated and KMT2A-rearranged cases and were further enriched for co-mutations commonly associated with juvenile myelomonocytic leukemia, such as PTPN11 (Figure 3g). This subtype was defined by differentiation arrest at an early monocytic stage (Figure 4a), displayed metabolically active transcriptional signatures (Figure 4b), and shared the immunophenotype with a recently described functionally defined monocytic stem cell population^42^ (Supplementary Figure 6h,i).

MR AML segregated into three distinct groups characterized by specific genomic (Figure 3g), biological (Figure 4b), and clinical features (Figure 3h,i). Stem-like MR AMLs showed minimal evidence of lineage priming, with the majority of pLSPCs residing in a stem-like state (Figure 4a). This subtype was enriched for complex karyotypes (Figure 3g) and exhibited strong TNF α- mediated inflammatory signaling (Figure 4b), likely reflecting its high degree of genomic instability. Notably, this archetype also included most MECOM-rearranged AML cases, which are well known for their stem-like properties^43^. In contrast, erythroid-primed MR AMLs were enriched specifically for monosomy 4, 9 or 22 (Figure 3g) and displayed lower levels of inflammatory signaling (Figure 4b,g). This archetype showed a higher proportion of residual healthy hematopoiesis (Supplementary Figure 6j), clear evidence of erythroid lineage priming (Figure 4a), and a more metabolically active phenotype (Figure 4b). Consistent with these biological characteristics, patients in this subgroup demonstrated improved responses to induction chemotherapy (Figure 3h) and more favorable outcomes following allogeneic stem cell transplantation compared with the stem-like MR AML archetype (Figure 4h). Notably, both stem-like and erythroid-primed MR AMLs were significantly enriched for p53 mutations (Figure 3g). Finally, pDC-primed MR AMLs were characterized by a highly immature cellular state with lineage priming toward early B cell and immature plasmacytoid dendritic cell (pDC) fates, accompanied by overproduction of mature pDCs (Figure 4a). This subtype was strongly enriched for RUNX1 mutations (Figure 3g) and associated with poor response to induction chemotherapy as well as markedly inferior overall survival (Figure 3h,i). These features closely align with the recently described pDC-AML subset^44^.

Together, these findings show that leukemic stem and progenitor cells within each AML archetype are arrested at unique differentiation states defined by recurrent genomic and epigenomic constraints, and delineate subtype-specific biological features, biomarkers, cellular hierarchies, and clinical characteristics.

### Genetics and differentiation state cooperatively drive response to induction chemotherapy

To elucidate the determinants of response to induction chemotherapy, we evaluated the explanatory power of individual feature sets using logistic regression and compared model performance using the Akaike Information Criterion (AIC), which balances goodness of fit and model complexity (Figure 5a; see methods). Therapy response was predominantly driven by leukemia-intrinsic features, which fell into three main categories: (i) genomic characteristics, including mutational profiles and established classification systems such as WHO and ELN22; (ii) measures of leukemic differentiation state, including the fraction of pLSCs, average pseudotime, and the LSC17 score; and (iii) cell cycle– and cellular activity–related properties, which had a more modest impact. In line with prior studies^19,31,45,46^, stem-like differentiation states, quiescent cell cycle activity, and adverse genetic subtypes were associated with inferior therapeutic response, whereas more differentiated and proliferative states, as well as favorable genetic profiles, correlated with better outcomes (Supplementary Figure 7a–c). Consistent with this, transcriptomic information from the broader pLSPC compartment provided the strongest association with induction response (Figure 5b), reflecting the previously described strong association of induction response with AML archetypes (compare Figure 3h). In contrast, information encoded in the immune environmental factors showed no substantial association with induction therapy response (Figure 5b, 2b).

**Figure 5.**
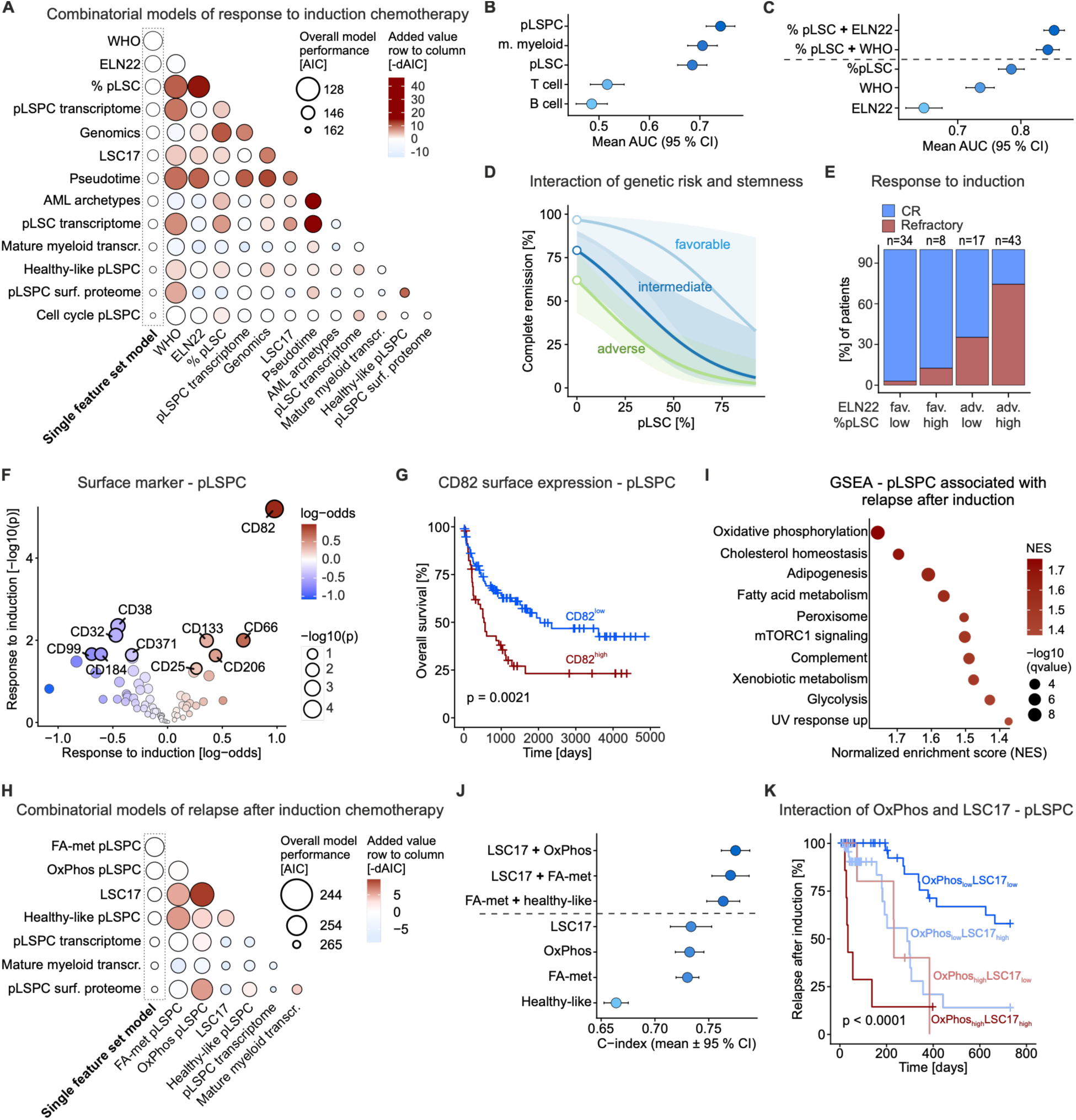
Combinatorial determinants of response and relapse following induction therapy. **(A)** Comparison of model performance for response to induction therapy across single feature set models and pairwise combinations, using selected features from Figure 2b. The leftmost column shows performance of individual feature sets, while the remaining columns represent pairwise feature combinations. Dot size reflects model performance, measured by the Akaike Information Criterion (AIC), and color indicates the incremental contribution of the row feature when combined with the column feature (−ΔAIC). Positive values denote improved model fit upon inclusion of the row variable. **(B)** Comparison of predictive performance across distinct cellular compartments for response to induction chemotherapy. Transcriptomes from each compartment were aggregated into patient-level pseudobulks and summarized using principal component analysis. Predictive performance was assessed using 5-fold cross-validation. Points represent mean area under the curve (AUC), and error bars indicate approximate 95 % confidence intervals (CI). **(C)** Predictive performance of models from selected individual or combined feature groups for response to induction chemotherapy. Performance was evaluated as described in (B). **(D–E)** Interaction of ELN22 genetic risk and pLSC burden. **(D)** Logistic regression–derived remission probabilities across pLSC levels, stratified by ELN22 risk. 95 % confidence intervals (CI) are indicated in shades. **(E)** Observed response rates stratified by ELN22 risk and pLSC burden (median split). **(F)** Volcano plot showing associations between surface marker expression in the pLSPC compartment and response to induction therapy. Surface markers were filtered to retain antibodies with Pearson correlation p < 0.05 between patient-level AbSeq abundance and matched RNA expression in pLSPCs, to reduce potential effects of nonspecific background or ambient signal. The x-axis shows the log-odds coefficient and the y-axis shows the −log10 p-value. Point color indicates effect direction and magnitude, and point size indicates significance. The top five markers by p-value are labeled for each direction of association. **(G)** Overall survival stratified by CD82 expression in the pLSPC compartment. **(H)** Comparison of model performance for relapse after induction therapy across single feature set models and pairwise combinations.. Model performance was evaluated using the same approach as in panel A. **(I)** Gene set enrichment analysis (GSEA) of pathways associated with relapse after induction. Genes were ranked by their association with relapse, and enrichment of Hallmark pathways was evaluated. The x-axis shows the normalized enrichment score (NES), with higher values indicating stronger positive association with relapse. Point size reflects statistical significance (−log10 q-value). **(J)** Predictive performance of individual and combined features for relapse after induction, evaluated by repeated cross-validation using the C-index. **(K)** Relapse after induction stratified by oxidative phosphorylation (OxPhos) gene set activity in pLSPC compartment and LSC17 expression. **(A–D, F)** Logistic regression was used for all models. **(H, J)** Cox proportional hazard models were used for all models. For all Kaplan–Meier analyses, high and low groups were defined using data-derived survival cutpoints selected to maximize separation of survival outcomes, and statistical significance was assessed using log-rank tests.

Although several of these features have been previously associated with clinical response, it remains unclear whether they act through shared or independent mechanisms. To quantify their combined contributions, we systematically evaluated pairwise combinations of relevant feature sets and compared their performance to the corresponding single feature set models (ΔAIC) (Figure 5a, see methods). Larger positive ΔAIC values indicate that the combined model better explains the outcome despite increased complexity, whereas negative values indicate that the additional variable increases complexity without sufficient improvement in fit. Despite partial redundancy, many pairwise combinations outperformed models based on single feature sets, indicating that variables retain independent, additive predictive value. In particular, differentiation-related features consistently added predictive power beyond current genetic classifications, and vice versa. For example, combining the fraction of stem-like leukemic cells (pLSCs) with WHO or ELN classification substantially improved prediction of induction therapy response (Figure 5c–e).

These findings were corroborated at the surface protein level (Figure 5f). Markers associated with myeloid maturation and reduced stemness (e.g., CD32, CD38, CD371) were enriched in leukemic stem and progenitor cells from patients with favorable responses, whereas stemness-associated markers (e.g., CD82 and CD133) predominated in poor responders (Figure 5f,g). A complete list of surface biomarker candidates associated with therapy response is presented in Supplementary Table 3.

Collectively, these findings show that both genetic and non-genetic leukemia-intrinsic features, particularly those linked to the leukemic differentiation state, independently and cooperatively influence the response to induction chemotherapy.

### Relapse dynamics after chemotherapy are driven by non-genetic leukemia-intrinsic programs

We next examined the biological determinants of disease relapse following induction therapy and how their interactions shape relapse risk. In contrast to the response to induction therapy, relapse dynamics were primarily driven by leukemia-intrinsic programs encoded in the pLSPC transcriptome and surface proteome that are independent of genetic drivers and differentiation state (Figure 2b). To identify such non-genotype associated biological processes, we performed GSEA in the pLSPC compartment using genes ranked by their predictive performance for time to relapse after induction chemotherapy therapy (Figure 5i).

This analysis revealed a strong enrichment of metabolic gene sets associated with early relapse, including oxidative phosphorylation (OxPhos), cholesterol homeostasis, and fatty acid metabolism (Figure 5i), all hallmark features of leukemic stem cell (LSC) metabolism ^47,48^. Importantly, these metabolic programs provided prognostic information beyond established leukemic stemness signatures and the above described healthy-like score, indicating that metabolic fitness contributes to early relapse independently of the underlying leukemic differentiation state (Figure 5h,j). Consistent with this, combinatorial stratification of patients based on OxPhos status and the LSC17 signature improved risk discrimination compared to either parameter alone, with patients exhibiting high OxPhos and high LSC17 scores relapsing rapidly after induction therapy, whereas those with low OxPhos and low LSC17 scores experienced considerably prolonged relapse-free survival (Figure 5k). Similarly, other metabolic programs, such as those related to fatty acid metabolism, considerably improved the risk stratification when combined with LSC17 or the healthy-like score (Figure 5h,j).

In line with its role as a marker of an immune-regulatory M2-polarized myeloid metabolic state characterized by enhanced oxidative phosphorylation, cholesterol homeostasis, and fatty acid metabolism^49^, high CD206 surface expression in leukemic stem and progenitor cells was associated with early relapse (Supplementary Figure 7d,e). A complete list of surface biomarker candidates associated with relapse risk after induction therapy is presented in Supplementary Table 3.

Collectively, these data suggest that relapse dynamics following induction chemotherapy is pre- determined at diagnosis by a combination of non-genetic, leukemia-intrinsic programs that, among others, relate to metabolic fitness of the leukemic stem and progenitor compartment.

### Healthy hematopoiesis and immune programming govern relapse dynamics post allogeneic stem cell transplantation

In contrast to the determinants of relapse following induction chemotherapy, processes associated with relapse after allogeneic stem cell transplantation (allo-SCT) showed less associations with leukemia-intrinsic factors and were instead dominated by leukemia-extrinsic regulatory mechanisms (Figure 2b). Among these factors, the cellular composition of the immune microenvironment at diagnosis emerged as a strong determinant of relapse dynamics following allo-SCT. In particular, high T cell frequencies at diagnosis were strongly associated with sustained remission post allo-SCT (Figure 6a-c). Within the T cell compartment, cytotoxic CD4 T cells and CD8 effector memory T cells re-expressing CD45RA (TEMRA) were particularly enriched in patients with durable responses, whereas naïve CD4 T cells were enriched in patients undergoing early relapse (Figure 6c). GSEA of genes associated with relapse after allo-SCT in the T cell compartment revealed that cytotoxic T cell programs were linked to long-term remission, whereas signatures indicative of naïve T cells were associated with earlier relapse (Figure 6a, Supplementary Figure 8a). In addition to T cells, signals encoded within the mature myeloid compartment, comprising monocyte and dendritic cell populations, were associated with relapse risk after allo-SCT in single feature set models (Figure 6a). Notably, gene programs associated with pro-inflammatory M1-like polarization, which support effective T cell priming^50^, were associated with long-term remission following allo-SCT (Figure 6a; Supplementary Figure 8b). Together, these findings suggest that a T cell–supportive immune microenvironment at diagnosis is associated with a favorable outcome post allo-SCT and may facilitate more effective graft-versus-leukemia responses following transplantation.

**Figure 6.**
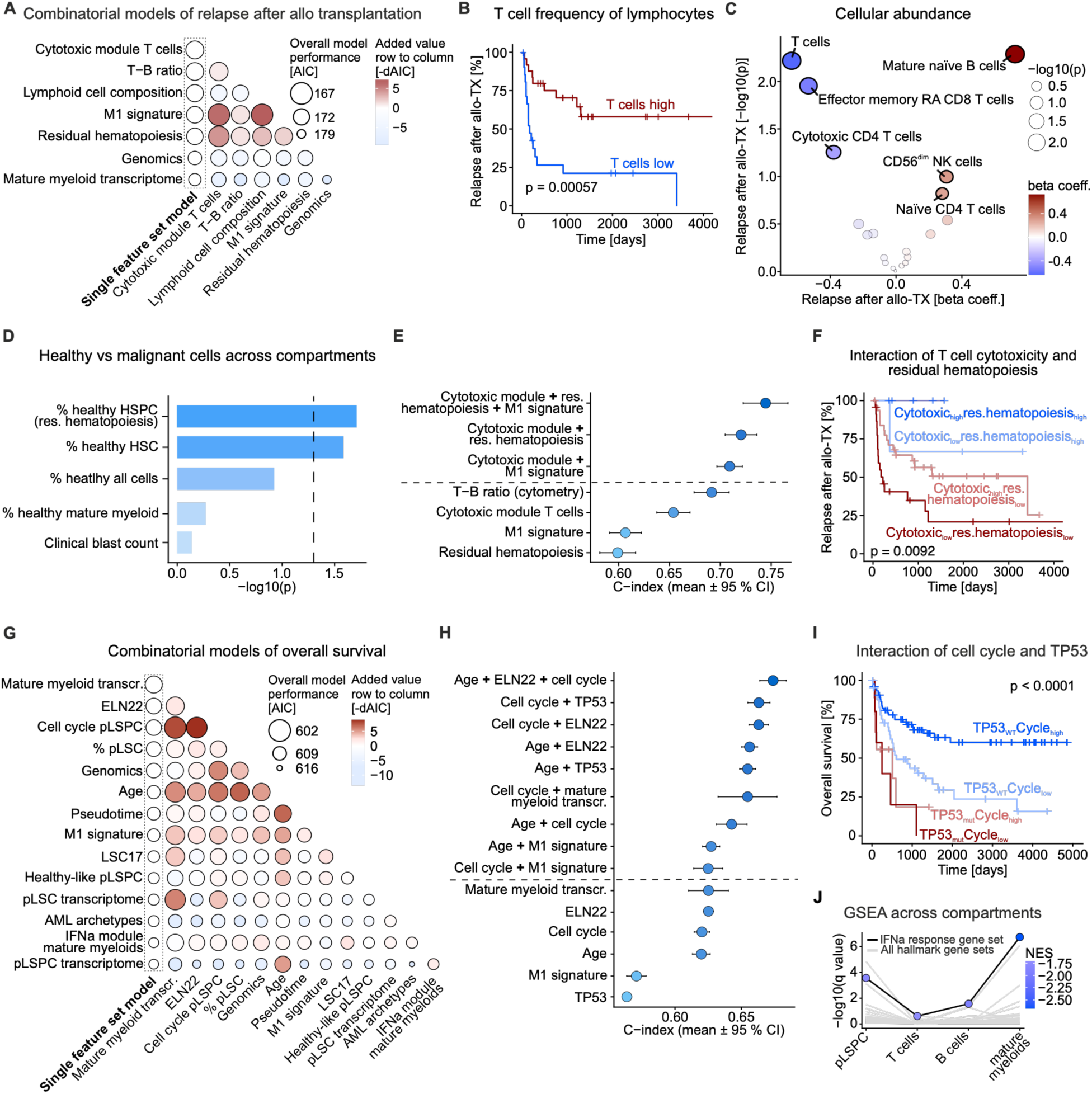
Combinatorial determinants of post-transplant relapse and overall survival. **(A)** Comparison of model performance for relapse after allogeneic transplantation across single feature set models and pairwise combinations, using selected features from Figure 2b, analogous to Figure 5a. Dot size reflects overall model performance using the Akaike Information Criterion (AIC), and color indicates the added value (−ΔAIC) of the row variable when added to the column variable. **(B)** Relapse after allogeneic transplantation stratified by T cell abundance derived from the flow cytometry measurements. **(C)** Volcano plot of associations between lymphoid cell subsets and relapse after allogeneic transplantation. Cox regression beta coefficients (x-axis) and statistical significance (y-axis; −log10 p-value) are shown; point color indicates effect direction and magnitude, and size reflects significance. **(D)** Association between predicted healthy cell fractions in different compartments and relapse after allogeneic stem cell transplantation (allo- TX). For each compartment, the fraction of cells predicted as healthy was tested as a predictor of relapse using Cox proportional hazards regression; clinical blast count was included for comparison. Bars show −log10 p-value from likelihood ratio tests against the null model. The dashed line indicates p=0.05. **(E)** Predictive performance of selected features and their combinations for relapse after allogeneic transplantation, evaluated by repeated cross-validation using the C-index (similar to Figure 5c). Points indicate mean area under the curve (AUC) and error bars represent approximate 95 % confidence intervals (CI) across folds and repeats. **(F)** Relapse after allogeneic stem cell transplantation (allo-TX) stratified by cytotoxic T cell activity and degree of residual hematopoiesis. **(G)** Comparison of model performance for overall survival across single feature set models and pairwise combinations. **(H)** Predictive performance of selected features and their combinations for overall survival, evaluated by repeated cross-validation using the C-index (similar to Figure 5c). **(I)** Overall survival stratified by cell cycle activity in pLSPCs and genomic TP53 status. **(J)** Gene set enrichment analyses (GSEA) across cellular compartments using genes ranked by their association with overall survival. Interferon alpha signaling is highlighted. The x-axis shows statistical significance (−log10 q-value), while point color represents the normalized enrichment score (NES), indicating direction and magnitude of enrichment. **(A, C, E, G, H)** Cox proportional hazard models were used for all models. For all Kaplan–Meier analyses, high and low groups were defined using data-derived survival cutpoints selected to maximize separation of survival outcomes, and statistical significance was assessed using log-rank tests. **(A,E,F)** T cell cytotoxic module gene set taken from Li et al.^59^

Notably, the extent of residual healthy hematopoiesis at diagnosis, defined as the fraction of healthy hematopoietic stem and progenitor cells relative to their malignant counterparts, was associated with longer remission following allo-SCT (Figure 6a; Supplementary Figure 8c). Similarly, the relative abundance of healthy HSCs compared to malignant pLSCs showed a comparable protective effect, whereas neither the overall ratio of healthy to malignant cells nor clinical blast counts exhibited a similar association (Figure 2b, Figure 6d). This suggests that the protective effect is specifically linked to the preservation of functional healthy stem and progenitor capacity, potentially reflecting a more intact bone marrow niche that supports efficient donor stem cell engraftment after transplantation.

In particular, the preservation of residual healthy hematopoiesis synergized with features of the immune microenvironment (Figure 6e), suggesting that these biological components contribute complementary information to the response to allo-SCT. Patients with both preserved healthy hematopoiesis and high T cell cytotoxicity exhibited the lowest risk of relapse following allo-SCT, whereas those with low residual hematopoiesis and reduced T cell cytotoxicity had the poorest outcomes (Figure 6f).

Collectively, these findings reveal a key role of leukemia-extrinsic factors and suggest that the immune microenvironment cooperates with the degree of residual healthy hematopoiesis at diagnosis to jointly shape relapse dynamics following allo-SCT.

### Combinatorial and overarching factors underlying overall survival

Conceptually, overall survival in AML reflects the cumulative impact of clinical events across a patient’s disease trajectory, together with additional overarching factors. Accordingly, overall survival was shaped by a complex interplay of the genetic and non-genetic determinants, substantially overlapping with those driving therapy response and relapse dynamics (Figure 6g, Figure 2a,b, Supplementary Figure 8d). In addition, several variables exerted largely independent effects on survival. As expected, patient age added prognostic information across most pairwise feature combinations (Figure 6g,h, Supplementary Figure 8e). Similarly, cell cycle activity in pLSPCs and genetic features contributed independent prognostic value when combined with other feature sets (Figure 6g,h). For example, cell cycle activity in pLSPCs interacted with TP53 mutational status or ELN22 classification to considerably influence overall survival (Figure 6h,i; Supplementary Figure 8f).

Notably, transcriptomic programs in mature myeloid cells emerged among the strongest predictors of overall survival in single feature set analyses and provided substantial additional value across several pairwise combinatory models (Figure 2b, 6g). GSEA of survival-associated genes in this compartment highlighted interferon response signaling as strongly linked to prolonged survival, consistent with its reported anti-leukemic effects in AML^51^ (Supplementary Figure 8g). Elevated type I interferon signaling was also associated with favorable prognosis across multiple cellular compartments, including both leukemic and microenvironmental cells (Figure 6j), suggesting a systemic effect that contributes to improved clinical outcomes.

Collectively, these findings suggest that overall survival emerges from coordinated interactions across biological layers and cellular compartments that drive individual outcomes, in conjunction with independent leukemia-intrinsic and systemic determinants.

## DISCUSSION

Clinical trajectories in cancer arise from the complex interplay between genomic alterations, cancer-intrinsic cellular processes, and the surrounding micro- and macroenvironment. While large-scale bulk multi-omics studies and single-cell technologies have independently transformed our understanding of tumor biology, they have largely examined these layers in isolation. Here, using AML as a paradigm, we combine large-scale single-cell multi-omics of a defined therapy-naïve cohort with a quantitative framework to systematically interrogate how interactions across biological layers and cellular compartments collectively shape clinical trajectories. Our findings provide several conceptual advances.

First, we demonstrate a strong link between genomic drivers and recurrent leukemia phenotypes within the broader stem and progenitor compartment, a relationship that becomes progressively obscured in more differentiated myeloid populations or when analyses are restricted to narrowly defined stem cell subsets. Building on this insight, we define AML archetypes as stable genotype– phenotype states that integrate genetic, epigenetic, transcriptional, and phenotypic features into coherent biological entities, thereby refining current classification systems and bridging genomic and differentiation-based frameworks. These archetypes show strong concordance with recent large-scale assemblies of bulk transcriptomic and epigenomic AML datasets^25,27,37^, while overcoming key limitations related to tumor purity, residual healthy hematopoietic cells, and microenvironmental impurities. While prior studies have established that leukemic cells occupy distinct positions along the hematopoietic hierarchy and that differentiation state shapes therapeutic response, our work provides a quantitative framework for these associations and reveals how genotype–phenotype relationships converge within the stem and progenitor compartment to drive distinct recurrent leukemic traits, biological programs, and clinical features.

Second, our study demonstrates that distinct clinical outcomes are governed by different combinations of biological layers and cellular compartments already present at diagnosis. Response to induction chemotherapy is primarily determined by leukemia-intrinsic features, particularly the interaction between genetic drivers and differentiation state. This is consistent with previous observations linking stem-like leukemic populations to therapy resistance^28,31^, but further shows that both effects provide independent information and can improve predictive performance when combined.

In contrast, relapse following induction therapy is driven predominantly by non-genetic leukemia-intrinsic programs, particularly those related to cellular metabolism. Emerging evidence has implicated oxidative phosphorylation and fatty acid metabolism in leukemic stem cell function and therapy resistance^47,52,53^. Our findings extend these observations by demonstrating that metabolic programs provide synergistic prognostic information independent of canonical stemness programs and genetic subtypes, suggesting that metabolic fitness represents a distinct layer of heterogeneity that contributes to shape relapse risk.

Finally, we identify a dominant role for several leukemia-extrinsic factors in shaping relapse dynamics following allogeneic stem cell transplantation. Specifically, the immune system at diagnosis, including a T cell supportive microenvironment, as well as the extent of residual healthy hematopoiesis, are key factors that synergize to determine subsequent post-transplant outcome. While these observations are consistent with a large body of studies emphasizing the role of immune context and T cell dysfunction during allogeneic stem cell transplantation^54–56^, our findings demonstrate that several of these features are already evident at initial diagnosis, thereby revealing a window of opportunity for improved patient stratification and early therapeutic modulation of the immune system prior to therapy.

Notably, the degree of residual healthy hematopoiesis provided complementary information to the immune landscape in stratifying relapse risk after allogeneic transplantation. This observation suggests that the preservation of a microenvironment supportive of hematopoiesis may be critical for effective donor stem cell engraftment and sustained leukemic control and is consistent with functional studies showing that efficient engraftment of healthy hematopoietic stem cells in immunocompromised mice is associated with improved outcomes^57,58^.

Together, these findings support a unifying model in which AML clinical trajectories are not determined by single biological factors but by specific combinations of mechanisms operating across biological scales and cellular compartments. Importantly, these combinations are outcome-specific: genetic and differentiation-related features dominate early treatment response, metabolic programs drive relapse after chemotherapy, and microenvironmental factors govern post-transplant outcomes.

Our findings have several important clinical implications. First, they indicate that AML risk stratification can be enhanced by integrating genetic with non-genetic and immune-based biomarkers. Second, they show that future disease trajectories are already encoded at diagnosis, opening opportunities for early intervention and more informed, personalized treatment decisions. Finally, we provide a catalogue of surface markers associated with distinct AML archetypes and clinical outcomes, facilitating rapid clinical translation.

In conclusion, we provide a comprehensive, multi-layered view of AML heterogeneity and establish a generalizable framework to resolve how biological layers interact to shape clinical outcomes. This work advances our understanding of AML at the systems level and lays the foundation for integrative approaches to precision medicine that account for the combinatorial nature of cancer biology.

## DATA & CODE AVAILABILITY

Code and processed data supporting the findings of this study are available for reviewer access at the following links: code repository https://github.com/agSHaas/200AML; processed data https://figshare.com/s/2b96974aa24617159a03

## ACKNOWLEDGEMENTS

We thank the NCT cell and liquid biobank for providing AML samples, the BIH Flow & Mass Cytometry Core Facility, the MDC Flow Cytometry Facility, the UKHD FACS Core Facility, the BIH/MDC Genomics Platforms, and the Core Unit Bioinformatics for technical support. This project was co-funded by the European Union (ERC, InteractOmics, 101078713 to S.H.). The views and opinions expressed are those of the authors only and do not necessarily reflect those of the European Union or the European Research Council; neither the European Union nor the granting authority can be held responsible for them. S.H., S.R., and L.V. received funding from the e:Med LeukoSyStem consortium (BMBF). S.H. and S.R. are supported by CRC1709 (DFG). S.H. also received funding from the Heisenberg Programme of the German Research Foundation (DFG), the e:Med LeukoSyStem consortium (BMBF), the TEP-CC consortium funded by the Bruno & Helene Jöster Foundation, the DKTK grant Identi-T, and CRC1588, CRC1444, TRR418, HA8790/3- 1, and the Germany’s Excellence Strategy – EXC 3118/1 – 533770413 (DFG). S.R. received funding from the Emmy Noether Programme of the DFG (RA 3166/1-1). L.V. was supported by the Fundación Asociación Española Contra el Cáncer (AECC laboratory grant). L.V. acknowledges support of the Spanish Ministry of Science and Innovation through the Centro de Excelencia Severo Ochoa (CEX2020-001049-S, MCIN/AEI /10.13039/501100011033), the Generalitat de Catalunya through the CERCA Programme and to the EMBL partnership. Schematics in Figure 1a were created with BioRender.

## METHODS

### Human samples

Bone marrow (BM) or peripheral blood (PB) samples from AML patients and healthy donors were obtained at Heidelberg University Hospital, and at Charité – Universitätsmedizin Berlin (Campus Virchow and Campus Benjamin Franklin), after informed written consent. BM aspirates were obtained from the iliac crest. Mononuclear cells from BM and PB were isolated by Ficoll density gradient centrifugation and cryopreserved in liquid nitrogen until further use.

All experiments involving human samples were approved by the ethics committees of Heidelberg University Hospital (S-112/2010, S-169/2017, S-672/2018, S-480/2011) and of the Charité – Universitätsmedizin Berlin (EA2/149/20, EA1/271/14, EA1/152/10, EA4/117/21) and were conducted in accordance with the Declaration of Helsinki.

### Sample processing and thawing

Primary patient samples were processed in two independent experimental cohorts in batches of 10 patients per day. For cohort integration, one healthy donor and one AML patient sample were used in both cohorts. Cryopreserved samples were rapidly thawed in a 37 °C water bath and immediately transferred into 20 mL RPMI medium supplemented with 10 % fetal calf serum (FCS), 0.5 mM EDTA, and 200 U DNase I, and centrifuged at 450 g for 5 min at 4 °C. Cell pellets were resuspended in 10 mL cold PBS supplemented with 2 % FCS (FB).

### Single-cell proteo-genomics

#### Cell sorting

Cells designated for single-cell proteo-genomics were centrifuged (450 g, 5 min, 4 °C) and resuspended in 250 µL PBS supplemented with 2 % FCS and 0.5 mM EDTA (FBE). Cells were stained with DAPI and Caspase 3/7 Green dye (Thermo Fisher Scientific; 1:2000) to assess cell viability. For each sample, 20,000 singlet DAPI^neg^Caspase^neg^ cells were sorted into low-binding 5 mL polypropylene tubes containing 400 µL FBE. For a subset of healthy control samples (n = 4), CD34 enrichment was performed to enhance resolution of hematopoietic stem and progenitor populations. For this purpose, cells were stained with a CD34-PE antibody (BD Biosciences, clone: 8G12) for 20 min, 4 °C in the dark. Following singlet and viability gating, cells were sorted into CD34^pos^ and CD34^neg^ fractions. Both fractions were processed independently, yielding paired datasets per control representing CD34-enriched and CD34-depleted compartments.

#### Sample multiplexing

Immediately after cell sorting, tubes were rinsed with 1 mL FBE and kept on ice. Cells were centrifuged (450 g, 5 min, 4 °C), resuspended in 20 µL sample multiplexing mix (BD Biosciences), and incubated on ice for 30 min. Sample multiplexing was performed using BD sample tags, with one unique tag assigned per patient to enable multiplexing on a single cartridge. In cohort 1, 1 µL of each tag was diluted in 19 µL FBE, whereas cohort 2 samples were labeled with 3 µL tag in 17 µL FBE to improve tag recovery. Subsequently, cells were washed three times with FBE and pooled during the final wash.

#### AbSeq labelling

For AbSeq surface labelling, the pooled suspension was incubated with 100 µL AbSeq antibody master mix for 30 min on ice (see Supplementary Table 2). After staining, cells were washed three times and resuspended in sample buffer (BD Rhapsody Cartridge reagent kit). Cell concentration was determined using Trypan Blue, targeting a loading of approximately 23,000 cells per BD Rhapsody cartridge.

#### Single-cell sequencing

Single-cell capture, cDNA synthesis, and library preparation were performed according to the manufacturer’s protocols of the BD Rhapsody™ Single-Cell Analysis System (“BD Rhapsody Express Single-Cell Analysis System” and “BD Rhapsody WTA, AbSeq, and Sample Tag Library Preparation Protocol”). Resulting libraries were quality checked by Qubit and Tapestation, pooled and sequenced using Illumina Novaseq S2 or S4 (Illumina; high-output mode). Ten BD Rhapsody libraries were generated for each of the two cohorts and sequenced in separate runs; cohort and sequencing-run identities were retained as annotations for downstream batch integration.

### High parametric flow cytometry

#### Experimental procedures and data acquisition

To comprehensively profile the cellular composition of the AML immune ecosystem, we employed three high-parameter panels^36^ designed to capture distinct aspects of the disease. (i) The “Identity panel” was developed to broadly define all major immune cell populations present in the AML microenvironment. (ii) The “T & NK cell panel” provided high-resolution characterization of T- and NK cell subsets. (iii) The “Myeloid panel” enabled in-depth analysis of myeloid differentiation states. Details of all panels, including antibody clones, fluorophore conjugates, and staining dilutions, are provided in Supplementary Table 2.

For high parametric flow cytometry, cells were distributed across up to three panels, depending on cell availability. Following centrifugation, cells were resuspended in 50 µL of the respective antibody master mix supplemented with eFluor 506 viability dye (1:1000, Thermo Fisher Scientific), Fc receptor blocking solution (1:20, BioLegend), and Brilliant Stain Buffer (10 uL per sample, BD Biosciences). Staining was performed for 20 min at 4 °C in the dark. Cells were subsequently washed with FB, centrifuged (450 g, 5 min, 4 °C), and resuspended in 200 µL FB for acquisition. Data was acquired using a BD FACSymphony A3 flow cytometer.

#### Analysis of flow cytometry data

For data analysis, FCS files were imported into FlowJo (BD Biosciences), and compensation was evaluated by inspection of N×N plots and adjusted where necessary. All parameters were visualized using a bi-exponential transformation, except for forward scatter (FSC) and side scatter (SSC), which were log-transformed. Axis limits were adjusted to facilitate visualization of gating boundaries.

Cell populations were identified by sequential gating, including exclusion of debris based on FSC and SSC, singlet discrimination using FSC area (FSC-A) and FSC wide (FSC-W), and removal of dead cells using an eFluor viability dye. Only live singlet cells were exported using FlowJo’s built-in export function with channel values preserved.

Exported CSV files were imported into R and analyzed using the Seurat v5 framework^60^. For the T cell and myeloid panels, up to 400,000 cells per sample were retained prior to FlowSOM^61^ clustering. Lineage-defining markers were used to restrict downstream analyses to the relevant compartment-specific clusters. For the Identity panel, up to 100,000 cells per sample were retained. For each flow cytometry panel, a Seurat object was then generated using BPCells^62^ to enable memory-efficient bit-packed data storage on a high-performance computing cluster. All measured marker expression values, as well as FSC-A and SSC-A, were included as features.

To correct for batch effects, Seurat’s sketch-based workflow was first applied to further downsample the dataset. A total of 50,000 cells per cohort were sketched, followed by dimensionality reduction using principal component analysis (PCA). Batch integration was then performed using anchor-based canonical correlation analysis (CCA), followed by construction of a shared nearest neighbor graph and Louvain clustering. As anchors, one healthy donor and one AML patient sample were used in both cohorts. These technical replicates were merged after batch integration. The full (unsketched) dataset was subsequently projected onto the integrated low-dimensional space using Seurat’s built-in functions, resulting in projected PCA and UMAP embeddings as well as cluster assignments. Notably, individual marker expression values were not batch-corrected. Cell type annotation was performed manually based on marker expression profiles. Following annotation, summary tables were generated to quantify cell abundances per patient. The human identity panel was used to derive an overview of major immune cell populations. To estimate lymphoid compartment composition, T cell abundances derived from the T cell panel were scaled by the overall T cell frequency obtained from the human identity panel.

### Variant calling using panel sequencing

Somatic mutations were assessed using DNA panel sequencing. For this purpose, genomic DNA was isolated from primary patient sample pellets (0.5 – 1 x 10^6^ cells) using the QIAamp DNA Mini Kit (Qiagen). A total of 100 ng DNA was used as input for library preparation with the TruSight Myeloid Sequencing Panel kit (Illumina) following the manufacturer’s instructions. Libraries were sequenced using 150-nucleotide paired-end reads with the NextSeq 1000/2000 P1 Reagent Kit (300 cycles) on a NextSeq 2000 platform (Illumina) or the HiSeq High Output Reagent Kit (300 cycles) on a HiSeq 2000 platform (Illumina). Sequencing and demultiplexing were performed by the EMBL Heidelberg Genomics Core Facility.

Demultiplexed FASTQ files were analyzed using the Illumina BaseSpace platform. Read alignment and somatic variant calling were performed using the TruSeq Amplicon App in BaseSpace with RefSeq hg19 annotation, a variant allele frequency (VAF) threshold of 2 %, and a minimum coverage of 20 reads, with read stitching enabled. For variant interpretation, a minimum VAF threshold of 5 % was applied, and synonymous single-nucleotide variants (SNVs) were excluded. Variants were classified according to the ACMG/AMP guidelines into Tier 1–5 variants^63^. Variants classified to Tier 1–3 were further manually curated and likely pathogenic variants were retained for downstream analyses. Patient samples were then classified into AML subtypes according to WHO 2022 classification incorporating cytogenetic features, gene fusions/rearrangements and mutation profiles^17^. In addition, samples were stratified prognostically based on the ELN 2022 criteria^19^.

### Pre-processing of single-cell proteo-genomic data

Raw sequencing data were processed using BD Rhapsody Sequence Analysis CWL pipeline (v.2.0b2) using GRCh38 as reference genome for alignment and GENECODE v29 for annotation. Low-quality cells were removed based on detected feature number (< 200), total counts (< 500), and mitochondrial transcript fraction (> 40 %).

Technical replicates were retained as separate entries for sequencing run–specific analyses and during batch correction and were subsequently combined for downstream analyses.

### Embedding and clustering of single-cell transcriptome

Single-cell RNA-seq data were processed in Seurat^60^. Cell-cycle phase was inferred after log-normalization using Seurat CellCycleScoring with default S-phase and G2/M gene sets. To prevent cell-cycle driven structure from driving the embedding and clustering, cells assigned to S or G2/M phase were initially excluded.

The remaining cells were split by sequencing run and independently normalized (log-normalization). Integration features were selected across runs (SelectIntegrationFeatures, 2,500 features), and each object was scaled and subjected to PCA. Datasets were integrated using reciprocal PCA anchor-based integration (RPCA using 30 dimension). UMAP embedding was generated using the first 60 principal components (PCs), and graph-based clustering was performed on the same dimensions using Louvain clustering at resolution 0.8.

S- and G2/M-phase cells were processed separately, split by cohort/day, integrated, and projected onto the non-cycling reference by Seurat’s MapQuery function to acquire UMAP and PCA coordinates, as well as cluster assignments.

Small clusters comprising < 1 % of all cells and dominated by a single sample (> 75 % of cluster cells) were merged with the most transcriptionally similar cluster. Similarity was defined by pairwise Pearson correlation of cluster-averaged gene expression profiles.

### Projection onto a healthy bone marrow reference

Cells were projected onto a healthy bone marrow reference atlas^35^ using scmap^64^, as implemented at: https://git.embl.org/triana/nrn/-/blob/master/Projection_Vignette/universal_ projection_function.R .

Normalized query and reference expression matrices were restricted to shared genes, and the ten nearest reference neighbors were identified for each query cell. Projected cell type was defined as the most frequent cell type among nearest neighbors, and lineage-specific pseudotime values were assigned by averaging nearest-neighbor pseudotime values from the reference across myelocytic, erythroid, megakaryocytic, B cell, and cDC trajectories.

### Malignant versus healthy cell classification

To restrict downstream analyses to malignant cells, we implemented a multi-step strategy to distinguish malignant from residual healthy cells. First, we defined high-confidence malignant cells using mutation- and copy-number-based evidence.

In samples carrying NPM1 mutations, malignant cells were identified by detection of at least one read containing the canonical 4-bp insertion at c.860_863 in single-cell RNA-seq data. Although coverage of the mutation-containing locus was sparse, mutation-positive cells were detected across most NPM1-mutated samples (Supplementary Figure 3b).

Next, large-scale chromosomal copy-number alterations were inferred from single-cell transcriptomes using inferCNV^65^. To increase specificity of the inferCNV calls, we evaluated a range of chromosome-span thresholds and selected a threshold of 0.4, as the minimum fraction of a chromosome required to be called gained or lost (Supplementary Figure 3d). Finally, cells were considered CNV-supported malignant only when the inferred CNV pattern was concordant with the clinical karyotype.

Together, NPM1 mutation detection and concordant CNV calls captured a broad range of malignant transcriptional states within the stem and progenitor compartment and constituted the reference malignant cells for the next steps (Supplementary Figure 3e).

### Malignancy estimation

To determine malignant status across all cells, including those from samples without detectable NPM1 mutations or informative chromosomal CNVs, we used a reference-based classifier. We constructed a reference set consisting of reference malignant cells, with evidence from NPM1 or CNVs, and reference healthy cells, using the healthy control samples (n=19,467 and n=11,332 cells respectively). For each cell, we identified its 100 nearest neighbors in the integrated PCA space. The number of healthy reference neighbors was used as a continuous predictor in a logistic regression model, with unsupervised cluster identity included as an additional covariate.

To avoid misclassifying malignant cells with healthy-like transcriptional states, we performed an additional sample-level enrichment correction. Among cells initially classified as healthy, we tested whether each unsupervised cluster was over-represented within a given AML sample relative to CD34 non-enriched healthy controls. Expected cluster frequencies were estimated from healthy controls, and enrichment was tested using a one-sided binomial test followed by Benjamini–Hochberg correction. Cells initially classified as healthy were reassigned as malignant when they belonged to non-lymphoid clusters significantly enriched in an AML sample after correction. Predefined lymphoid or mature immune clusters were excluded from this reassignment step.

### Validation of malignant-cell classification in an external ground-truth cohort

We validated this malignancy-classification framework using the CloneTracer cohort^32^, which provides independent ground-truth malignant and healthy annotations. CloneTracer cells were projected into the same reference space and classified using the same nearest-neighbor and logistic-regression strategy, omitting the enrichment correction because the cohort was CD34-enriched. Predicted labels were compared with CloneTracer ground truth across all cells and within the hematopoietic stem and progenitor cell compartment.

### Patient-level pseudobulk generation

Based on healthy bone marrow projection and malignancy estimation, cells were classified as putative leukemic stem cells (pLSCs), putative leukemic stem and progenitor cells (pLSPCs), mature myeloid cells, lymphoid cells, T cells, and B cells. pLSCs were defined as malignant cells projected to the healthy bone marrow “HSCs & MPPs” reference label, whereas pLSPCs were defined more broadly as malignant cells projected to stem and progenitor reference populations. Patient-level pseudobulks were generated separately for each of these six compartments by aggregating RNA counts by patient. Genes were retained if detected in more than five samples and if total expression exceeded the median total expression across genes. Pseudobulk matrices were log-normalized and run-related batch effects were corrected using ComBat^66^. For dimensionality reduction, the 1,000 most variable genes were selected based on dispersion, and PCA was performed on the normalized, batch-corrected expression matrix.

### Quantification of gene-set scores

Gene-set scores were computed at single-cell level and, where appropriate, averaged per patient within the relevant compartment. The LSC17 score was calculated as a weighted expression score using the canonical 17-gene LSC17 signature and published gene weights^31^. Specifically, scaled expression values for the LSC17 genes were multiplied by their corresponding weights and summed per cell to generate a per-cell LSC17 score.

Other transcriptional programs, including cell-cycle activity, lineage programs, and custom pathway scores, were quantified using Seurat AddModuleScore. All gene sets used for score calculation are listed in Supplementary Table 4.

### Definition of feature sets for cross-scale association analyses

For comprehensive analyses of clinical, genetic, transcriptomic, proteomic, microenvironmental, and cell-biological features, their relationships with each other, and their associations with clinical outcomes (Figures 2, 5, and 6), feature sets were defined as follows.

Clinical features included continuous variables, such as age and clinical blast count, and categorical variables, including WHO 2022 classification and ELN 2022 genetic risk.

Genetic features were derived from targeted sequencing and cytogenetic annotations. Features with more than 35 % missing values or fewer than five positive cases were excluded. Missing values in the remaining genetic features were imputed using multiple imputation by predictive mean matching with the mice package. The imputed genetic feature matrix, including cytogenetic group annotation, was then reduced by multiple correspondence analysis (MCA), and the first five resulting components were used as the genomic feature set.

For compartment-specific transcriptomic feature sets, the first 15 PCs derived from patient-level pseudobulks were used.

For the pLSPC surface proteome feature set, AbSeq counts were aggregated by patient and log-normalized. Antibody features corresponding to HLA-DR and TCR probes were excluded to avoid clone-specific effects and reduce batch effects. PCA was then performed on the normalized surface-marker pseudobulk matrix, and the resulting 15 first components were used as the pLSPC surface-proteome feature set.

Lymphoid microenvironment composition was derived from annotated flow cytometry data. T cell and B cell subset abundances were extracted, zero values were replaced using count zero multiplicative replacement implemented in R package zCompositions^67^, and compositional values were transformed using a centered log-ratio transformation. Features were scaled, and PCA was performed; the resulting first five components were used to represent lymphoid microenvironment composition.

Pseudotime features were derived from projection onto the healthy bone marrow reference. Lineage-specific pseudotime values for myelocytic, erythroid, megakaryocytic, B cell , and cDC trajectories were converted to percentile ranks from 0 to 1. A combined pseudotime score was calculated per cell from the maximum percentile-scaled lineage pseudotime across trajectories and then percentile-scaled again. Patient-level pseudotime was obtained by averaging this combined score across malignant cells.

The percentage of pLSCs was defined as the proportion of malignant cells in each sample assigned to the projected “HSCs & MPPs” reference label. Residual hematopoiesis was defined as the proportion of cells classified as healthy within the pLSPC compartment. Cell-cycle activity was quantified using the S-phase gene set averaged per patient across pLSPCs.

A healthy-like transcriptional score was derived from differential expression between AML and healthy progenitor-cell pseudobulks. For AML samples, pseudobulks were generated from pLSPC compartment; for healthy controls, pseudobulks were generated from stem and progenitor cells. To account for differences in composition, projected cell types were collapsed into five major groups: HSC/MPP, B/pDC progenitor, lymphomyeloid progenitor, promyelocyte, and megakaryocyte/erythroid progenitor. These compositional covariates were centered log-ratio transformed and included in the DESeq2^68^ model together with batch and AML-versus-healthy status. Differential expression was estimated with DESeq2, and healthy-associated genes were defined as genes with adjusted p < 0.05 and log2 fold change < −1.5 in AML versus healthy controls. These genes were used to compute per-cell healthy-like scores and averaged across the pLSPC compartment for each sample.

### Cross-scale association analysis

We performed reciprocal prediction analyses across patient-level variables and feature sets (Figure 2a). For each ordered feature-pair comparison, the row feature set was treated as the outcome and the column feature set as the predictor. Healthy controls were excluded from this analysis.

The model type and summary statistic were chosen according to the outcome type. For single continuous outcomes, such as pseudotime or LSC17 score, we used linear regression and quantified association strength using adjusted R^2^. For categorical outcomes, such as WHO 2022 subtype or ELN 2022 risk group, we used multinomial regression and quantified model fit using adjusted McFadden’s pseudo-R^2 69^. McFadden’s pseudo-R^2^ was defined as R^2^_McFadden = 1 − logL_full / logL_null, where logL_full and logL_null are the log-likelihoods of the full and null models, respectively. The adjusted version was calculated as R^2^_McFadden,adj = 1 − (logL_full − k) / logL_null, where k is the number of estimated parameters in the full model.

For multidimensional numeric feature sets, such as PCs representing compartment-level transcriptomes or the surface proteome, we used multivariate linear models. Association strength was summarized using an adjusted trace-based multivariate R^2^, calculated from the reduction in residual variation across all outcome dimensions. The unadjusted trace-based R^2 70^ was defined as R^2^_trace = 1 − tr(E_full) / tr(T), where E_full is the residual sum-of-squares-and-cross-products matrix from the full model, T is the total sum-of-squares-and-cross-products matrix of the outcome feature set, and tr(.) denotes the matrix trace. The adjusted version was calculated as R^2^_trace,adj = 1 − (1 − R^2^_trace) × (n − 1) / (n − k), where n is the number of samples and k is the number of estimated parameters in the full model, including the intercept.

For each comparison, the full model was compared with an intercept-only null model. Statistical significance was assessed using F-tests for continuous single-feature outcomes, likelihood ratio tests for categorical outcomes, and Pillai’s trace test for multivariate numeric outcomes. P-values were adjusted across comparisons using the Benjamini–Hochberg method. Associations were visualized when the adjusted p-value was < 0.1 and R² type metric was > 0.05.

For clustering of variables in the association map, pairwise reciprocal association strengths were determined as the minimum of the two R²-based scores, and distance was defined as one minus this reciprocal score. Hierarchical clustering using Ward’s method was then performed.

### Clinical endpoints

Outcome analyses were restricted to AML patients treated with intensive chemotherapy. Healthy controls, patients with acute promyelocytic leukemia, and patients receiving non-intensive treatment were excluded unless stated otherwise.

**Response to induction therapy** was binarized according to best response to first-line chemotherapy as complete remission (CR) versus refractory disease. CR was defined by bone marrow blasts < 5 % in a non-aplastic bone marrow, absence of circulating blasts and absence of extra-medullary disease. If not explicitly stated in the bone marrow report, CR can be inferred from MRD negativity or documented remission in fluorescence in situ hybridization (FISH) or flow cytometry reports. Failure to achieve remission, progression or early discontinuation of therapy was marked as refractory disease.

**Overall survival** was defined as the time from diagnosis to death from any cause, with surviving patients censored at last follow-up.

To separately evaluate relapse dynamics before and after allogeneic transplantation, two relapse endpoints were defined.

**Relapse after induction** was measured from achievement of remission to the earliest of relapse, death, allogeneic stem cell transplantation, or last hematologic assessment. Relapse was counted as the event of interest; death without relapse, transplantation before relapse, and loss to follow-up were treated as censoring events. Patients for whom transplantation occurred before documented remission were excluded from this endpoint.

**Relapse after allogeneic stem cell transplantation** was evaluated only in patients who underwent transplantation before relapse. Time was measured from transplantation to relapse, death, or last hematologic assessment. Relapse was counted as the event of interest, whereas death without relapse and last assessment were treated as censoring events.

The number of evaluated patients was n = 134 for response to induction, n = 141 for overall survival, n = 113 for relapse after induction, and n = 63 for relapse after allogeneic transplantation.

### Feature-set associations with clinical endpoints

Associations between patient-level feature sets and clinical endpoints were evaluated by fitting one model per feature set and endpoint. Feature sets shown in the rows of Figure 2b were used as predictors. Response to induction therapy was modeled using logistic regression, whereas overall survival, relapse after induction, and relapse after allogeneic stem cell transplantation were modeled using Cox proportional hazards regression.

For each endpoint, the full model containing the feature set was compared with an intercept-only null model using a likelihood ratio test. Model improvement was additionally summarized by ΔAIC, calculated as AIC (null model) – AIC (full model), so that positive values indicate improved fit after accounting for model complexity. For Cox models, proportional hazards assumptions were evaluated using Schoenfeld residuals^71^. When significant proportional hazards violations were detected for predictors that were also nominally associated with outcome, time-varying effects were modeled using penalized spline terms and retained if significantly associated with the outcome.

P-values were adjusted across all tested models using the Benjamini–Hochberg method. Associations were displayed when adjusted p < 0.1 and ΔAIC > 0.

### Embedding and clustering of AML archetypes

Pseudobulks for pLSPC compartment were generated as described above and the first 15 PCs were selected based on evaluation of clustering stability across different PC ranges using silhouette scores. A nearest-neighbor graph was constructed using cosine distance, 15 nearest neighbors, and the selected PCs. Leiden clustering was performed on this graph at resolution 0.7, selected based on silhouette scores computed across multiple random seeds. UMAP visualization was computed from the same nearest-neighbor graph.

### Archetype prediction in external bulk RNA-seq datasets

To link AML archetypes to additional modalities and increase patient numbers, we inferred archetype assignments in the bulk RNA-seq cohort from the TCGA dataset^12^ and data from Severens et al.^37^.

### AML Cell state deconvolution

We first estimated the abundance of AML cell states, defined from our single-cell reference map, in bulk RNA-seq samples using CIBERSORT^72^. The signature matrix was generated from the single-cell dataset by randomly sampling up to 200 cells per merged Louvain cluster, normalizing expression to CPM, and restricting the matrix to the 5,000 most variable genes shared between the single-cell and external bulk RNA-seq datasets. The resulting signature matrix was used to deconvolve CPM-normalized bulk RNA-seq profiles and estimate cell-state composition in each external sample.

### Archetype signatures

We also generated archetype-specific transcriptional signatures from the pLSPC pseudobulks. For each archetype, DESeq2 was used in a one-versus-rest framework with batch as a covariate. Genes were ranked by the Wald statistic, and archetype-enriched and archetype-depleted gene sets were defined using adjusted p-value, shrinkage-adjusted log2 fold change, and minimum expression thresholds. Gene sets were restricted to a maximum of 200 genes, and genes appearing in more than two archetype-up signatures were removed to improve specificity. Single-sample enrichment scores for these archetype signatures were then computed in both internal and external datasets using ssGSEA.

### Archetype prediction

was performed by combining ssGSEA-derived archetype signature scores with CIBERSORT-derived cell-state composition estimates. Features were normalized within datasets and across samples before classification. A class-weighted random forest classifier was trained on the internal cohort using known archetype labels. Feature importance was estimated from the full model, and the number of features retained for prediction was selected by evaluating out-of-bag error across a grid of top-ranked feature counts. The final model with 17 features was applied to the external bulk RNA-seq cohorts to assign archetype probabilities and predicted archetype labels.

Baseline confidence thresholds were selected from internal out-of-bag predictions to maintain ≥ 90 % accuracy among retained calls. Since HOX-stem archetype (0) was the largest subgroup and could otherwise accumulate more false-positive assignments by sample number, a stricter threshold was applied for this class (+ 0.10 top-class probability and + 0.05 probability margin). Samples with ambiguous archetype predictions were excluded from downstream analyses.

### DNA methylation data preprocessing and analysis

TCGA-LAML Illumina HumanMethylation450 array data^12^ were obtained as beta values. Probes with missing values in any sample were removed. To reduce technical and genetic confounding, probes mapping to sex chromosomes, probes overlapping common SNPs, and previously reported cross-reactive probes were excluded. Beta values were constrained to the interval [10– 6, 1 – 10 – 6] and converted to M-values using M = log2(beta / (1 – beta)).

For lineage-priming methylation analyses, monocyte- and granulocyte-associated methylation loci were defined using the FlowSorted.Blood.450k reference dataset^73^. Raw reference methylation data were preprocessed using noob normalization, and differential methylation between monocyte and granulocyte samples was tested on M-values using limma. Marker loci were retained if they showed adjusted p-value ≤ 1 × 10⁻⁶ and absolute delta beta ≥ 0.20. Enriched loci were then intersected with probes retained in the processed TCGA-LAML methylation matrix. To score granulocyte-biased methylation in TCGA AML samples, M-values for retained granulocyte-enriched loci were z-scored across samples for each probe and averaged per sample.

### Prediction of WHO subtype from compartment-specific pseudobulk clustering

Pseudobulks and principal component decomposition were generated as described above. For each compartment, a nearest-neighbor graph was constructed from the first 15 PCs using cosine distance. Leiden clustering resolution was selected separately for each compartment by screening candidate resolutions across multiple random seeds and evaluating silhouette scores. To assess the predictive utility of compartment-specific transcriptomic structure, clustering labels from each compartment were used as predictors of WHO 2022 subtype in multinomial regression models. Model performance was evaluated using 5-fold cross-validation, and mean classification accuracy across folds was reported (Figure 3a).

### Conservation of archetype-specific structure across data layers

To evaluate whether AML archetypes were reflected across independent data layers, we generated unsupervised cluster labels separately for methylation, regulon activity, surface proteome and lymphoid composition. For each data-layer we proceeded as follows. PCA was first performed, and the number of PCs used for graph construction was selected by inspecting cumulative explained variance and selecting an approximate elbow point. Nearest-neighbor graphs were then constructed in the selected PCA space, and Leiden clustering was performed. Clustering resolution was selected by screening resolutions across multiple random seeds and evaluating silhouette scores.

For methylation data, the 10,000 most variable CpG methylation probes were selected by median absolute deviation of M-values across samples and used for PCA-based embedding and clustering. For surface proteome data, patient-level pLSPC AbSeq pseudobulks were processed as described above. SCENIC ^38^ was applied to single-cell pLSPCs to infer regulon activity, and the resulting regulon-by-cell matrix was used for downstream steps. Lymphoid composition clusters were generated from flow cytometry-derived lymphoid composition as described above.

To quantify overlap between layer-specific cluster labels and AML archetypes, we calculated the adjusted Rand index between each layer-specific clustering and the AML archetype labels. For TCGA methylation data, archetype labels were taken from bulk RNA seq–based archetype predictions.

### Differentiation trajectory visualization

Differentiation trajectories were visualized in semi-circular coordinates for malignant pLSPCs (Figure 4a). The radial coordinate represented combined pseudotime, and the angular coordinate represented lineage bias. Lineage bias was inferred from module scores for seven lineages: megakaryocytic, erythroid, eosinophil–basophil, neutrophilic, promonocytic, pDC, and B cell. Lineage marker sets were derived from published data^29,35^ and refined using marker expression in healthy reference progenitor populations. Per-cell lineage scores were converted to softmax weights, and the final angle was determined by the weighted sum of unit lineage vectors. Patient points indicate the mean pLSPC position per archetype. Arrows indicate lineage directions with mean softmax weight > 0.2; arrow length reflects the 80^th^ percentile of pseudotime among cells assigned to that dominant lineage.

Detailed implementation, including exact parameter settings and intermediate processing steps, is provided in the accompanying analysis code (see code availability).

### Detection of outcome-associated pathways by GSEA

To identify transcriptional programs associated with clinical outcomes (Figures 5i, 6j; Supplementary Figures 7c, 8g), we used the compartment-specific patient-level pseudobulks described above. For each compartment and clinical endpoint, genes were tested individually for association with outcome. Response to induction was modeled using logistic regression, whereas overall survival, relapse after induction, and relapse after allogeneic transplantation were modeled using Cox proportional hazards regression. Proportional hazards assumptions were evaluated using Schoenfeld residuals. If a gene had an unadjusted Cox model p < 0.05 and showed evidence of proportional-hazards violation, a time-varying Cox model with penalized spline terms was fitted, and the likelihood-ratio-test p-value for the fitted model was used for ranking. Otherwise, the Wald p-value for the gene coefficient from the standard Cox model was used. Genes were ranked using a direction-aware significance score, defined as −log10(p) × sign(β), where p is the outcome-specific gene-level p-value and β is the logistic-regression or Cox-regression coefficient. Ranked gene lists were used for GSEA with gene sets from MSigDB Hallmarks together with selected custom immune programs. Gene sets containing fewer than 5 or more than 300 genes were excluded.

### Association of pairwise feature-set combinations with clinical endpoints

To evaluate whether selected feature sets contributed overlapping or complementary information for clinical outcome prediction (Figures 5a,h and 6a,g), we compared models based on individual feature sets with models based on pairwise feature-set combinations. For each outcome, analyses were restricted to complete cases across the selected feature sets.

Response to induction was modeled using logistic regression. Overall survival and relapse endpoints were modeled using Cox proportional hazards regression, as described above. A model was fit for each feature set and each pairwise combination. Model performance was summarized using AIC, with lower AIC indicating better fit after penalizing for model complexity. Added value was quantified by ΔAIC comparing the combined model with the corresponding single feature set model. Specifically, ΔAIC was defined as AIC (single feature set model) – AIC (combined model), so that positive values indicate improved model fit after adding the second feature set despite increased complexity.

Added value was evaluated in both directions depending on which feature set is treated as the baseline model. In displayed graphs, only one triangular half of the matrix is displayed to emphasize cases in which generally weaker predictors improve stronger baseline models.

### Cross-validated evaluation of selected prediction models

Feature sets and their combinations prioritized by prior analyses were evaluated by 10-times repeated 5-fold cross-validation to compare their internal predictive performance and relative contribution to clinical outcome models (Figure 5b,j, 6e,h). For each repeat, folds were generated to preserve the outcome distribution: response class for induction response models and event status for survival models.

Response to induction was modeled using binary logistic regression, and performance was assessed from held-out predicted probabilities using AUC. Relapse and survival endpoints were modeled using Cox proportional hazards regression, and performance was assessed on held-out folds using Harrell’s C-index.

Metrics were averaged across folds and repeats. Approximate 95 % confidence intervals were calculated as mean ± 1.96 standard errors across repeated cross-validation results.

### Kaplan–Meier stratification of continuous predictors

For Kaplan–Meier visualization, continuous predictors were dichotomized using R function survminer::surv_cutpoint^74^, which identifies data-derived cutpoints based on maximally selected rank statistics. For analyses combining two continuous predictors, the dichotomized variables were crossed to define four groups.

---

**Supplementary Figure 1.**
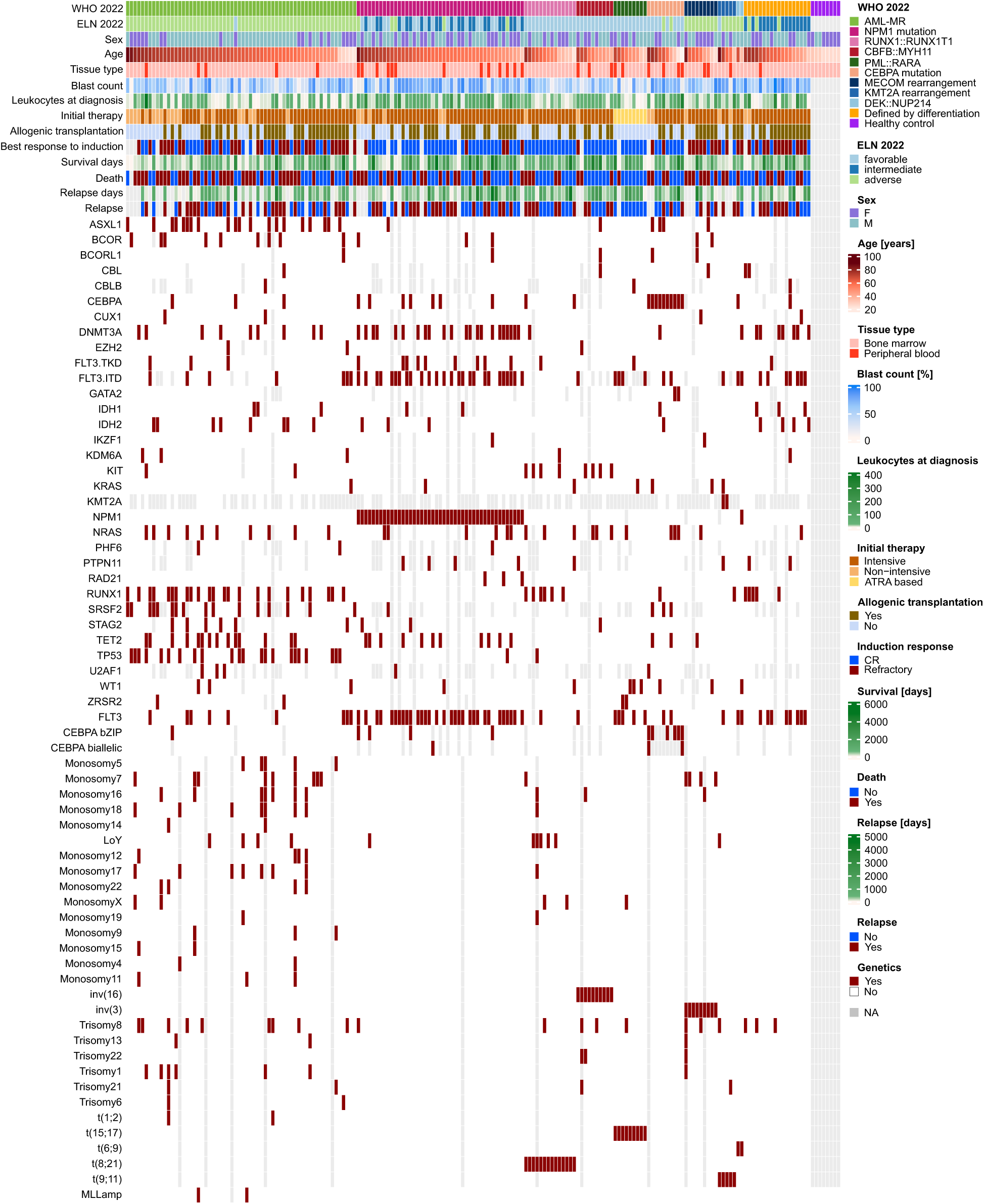
Integrated overview of patient metadata and genomic landscape. Columns correspond to individual patients. Top tracks show clinical, demographic, and treatment variables, while the bottom tracks depict recurrent mutations and chromosomal abnormalities.

**Supplementary Figure 2.**
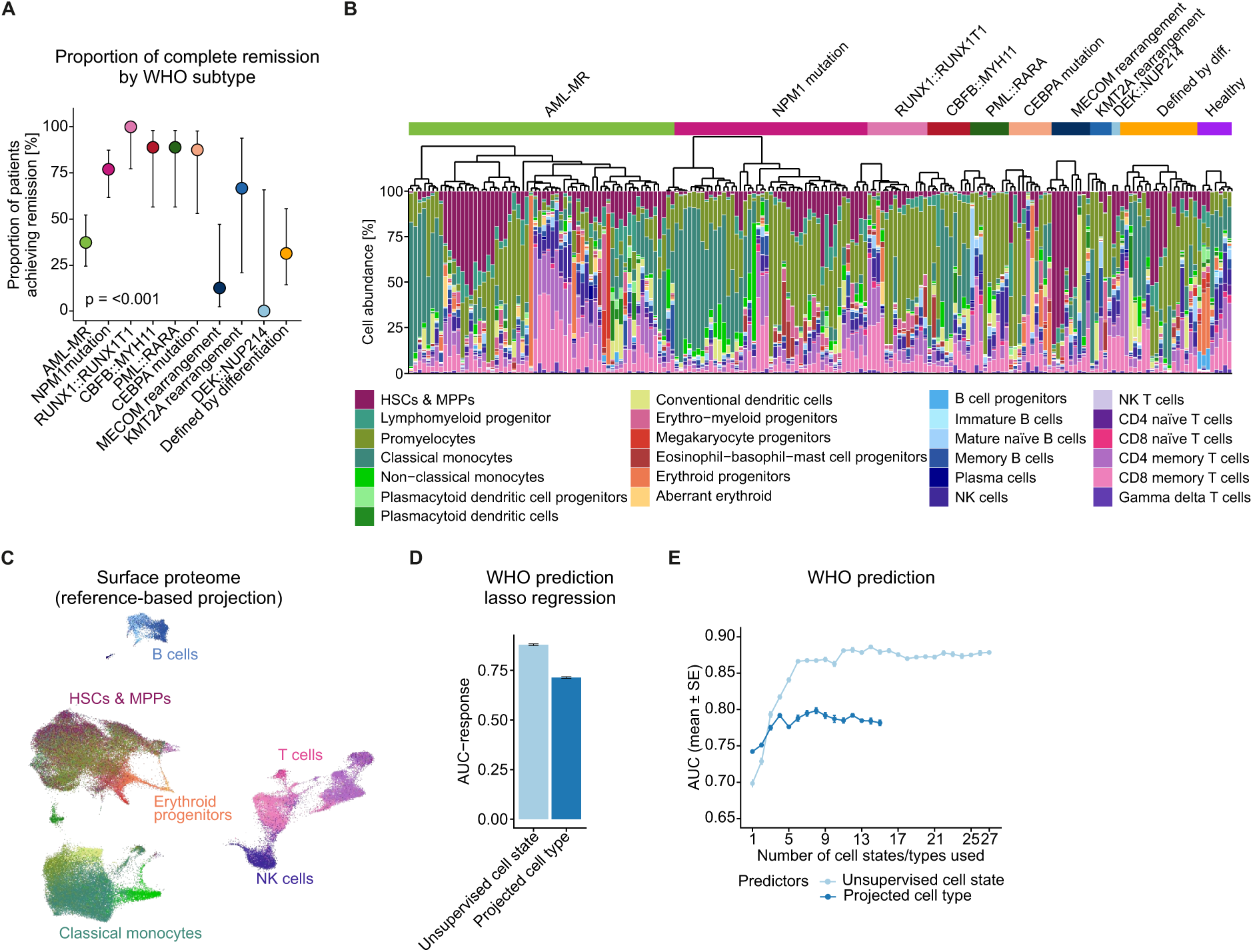
Clinical and cellular features stratified by genetic subtypes. **(A)** Response to induction therapy across WHO subtypes. Percentage of patients achieving complete remission is indicated, with error bars indicating 95 % confidence intervals calculated using the Wilson method. Overall differences across archetypes were assessed using Fisher’s exact test based on contingency tables of response categories. **(B)** Cell type composition of patient samples based on single-cell transcriptomic layer of single-cell proteo- genomic data, inferred by projection onto a healthy bone marrow reference^1^. Samples are shown as stacked proportions. Hierarchical clustering was performed within WHO subtypes. **(C)** UMAP representation of the surface proteome layer from the single-cell proteogenomic dataset. AbSeq counts were log-normalized, and HLA-DR and TCR probes were excluded before integration. Samples were split by cohort/run and integrated using Seurat anchor-based integration with the first 40 principal components (PCs). UMAP was generated from the integrated surface proteome representation using the first 40 PCs. Cells are colored by reference-projected cell type labels. **(D,E)** Prediction of WHO 2022 subtype from sample-level cell state (by unsupervised clustering) and projected cell type abundances. **(D)** Multinomial lasso models were evaluated by repeated stratified cross-validation; bars show mean one-vs-rest multiclass area under the curve (AUC), with black error bars indicating standard error. **(E)** Multinomial models were fit using increasing numbers of predictors (cell types and cell states), ordered by mean abundance from most to least frequent. Lines show AUC and error bars indicate standard error (SE) across repeated 5-fold cross-validation.

**Supplementary Figure 3.**
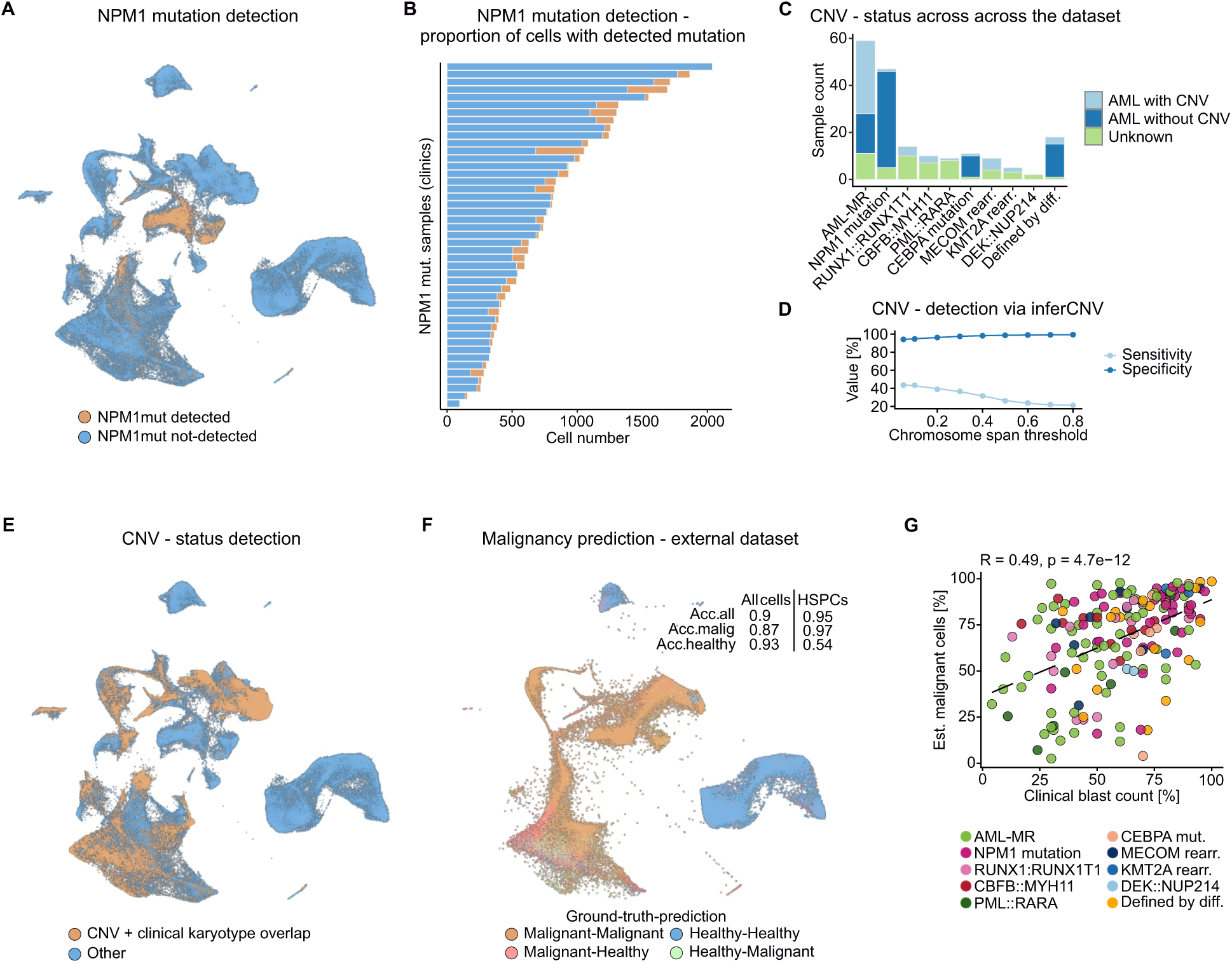
Identification and prediction of healthy versus malignant cells. **(A)** Detection of the canonical NPM1 4-bp TCTG insertion (c.860_863) in single-cell RNA-seq data, visualized on UMAP; mutation-positive cells are indicated. **(B)** Fraction of cells with detected NPM1 mutation per sample within the leukemic stem and progenitor compartment of NPM1 mutant patients. **(C)** Sample-level copy number variation (CNV) status stratified by WHO subtypes. CNVs were defined based on clinically detected monosomies and trisomies from karyotyping. **(D)** Sensitivity and specificity of inferCNV-based^2^ CNV detection across chromosome-span thresholds for monosomies and trisomies. Clinical karyotyping was used as the reference, and thresholds indicate the minimum chromosome fraction called as gained or lost. **(E)** Cells with concordant CNV calls between inferCNV (0.4 chromosome-span threshold) and clinical karyotyping are shown; these represent malignant cells supported by both approaches. **(F)** External validation of malignancy prediction using the CloneTracer dataset^3^. Cells were projected onto the reference space and classified as malignant or healthy using the same prediction framework as for the primary dataset. CloneTracer cells were mapped onto the reference UMAP embedding and colored by ground truth–prediction concordance. Accuracy is reported for all cells (Acc. all) and for the hematopoietic stem and progenitor cell compartment (HSPC). Acc. malig. and acc. healthy denote class-specific accuracies for ground-truth malignant and healthy cells, respectively. **(G)** Correlation between clinical blast count and estimated % of malignant cells. Spearman’s rank correlation coefficient is indicated. Each point represents an individual patient sample, colored by WHO 2022 classification. Dashed lines show ordinary least squares fits for visualization.

**Supplementary Figure 4.**
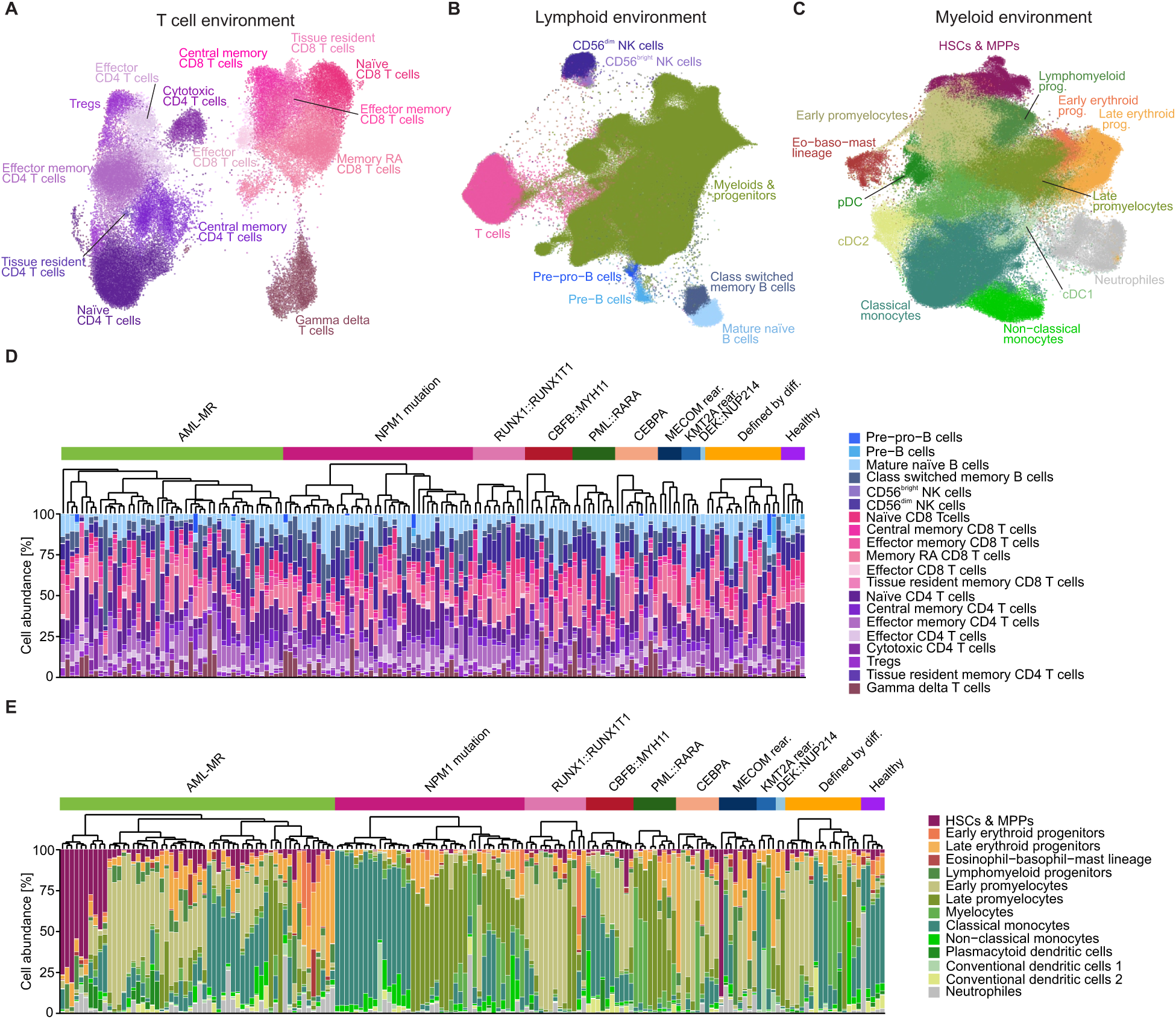
Mapping of the immune microenvironment using ultra-high plex flow cytometry. (A–C) UMAP embeddings and cell type annotation based on three flow cytometry panels^4^. For each panel, the same cell number per patient is visualized: **(A)** T cell– focused panel (n=7,299,556 cells analyzed, n=66,640 cells visualized), **(B)** broad immune panel (n=17,077,819 cells analyzed, n=142,082 cells visualized), and **(C)** myeloid-focused panel (n=17,190,943 cells analyzed, n=395,296 cells visualized). **(D)** Composition of the lymphoid microenvironment. NK and B cell subset proportions are quantified using the broad immune panel. T cell subtype frequencies were determined using the T cell–focused panel and scaled to the total T cell abundance derived from the broad immune panel. **(E)** Composition of the myeloid compartment based on the myeloid-focused panel. **(D,E)** Cell type composition is shown per sample as stacked proportions; samples are hierarchically clustered within WHO subtypes.

**Supplementary Figure 5.**
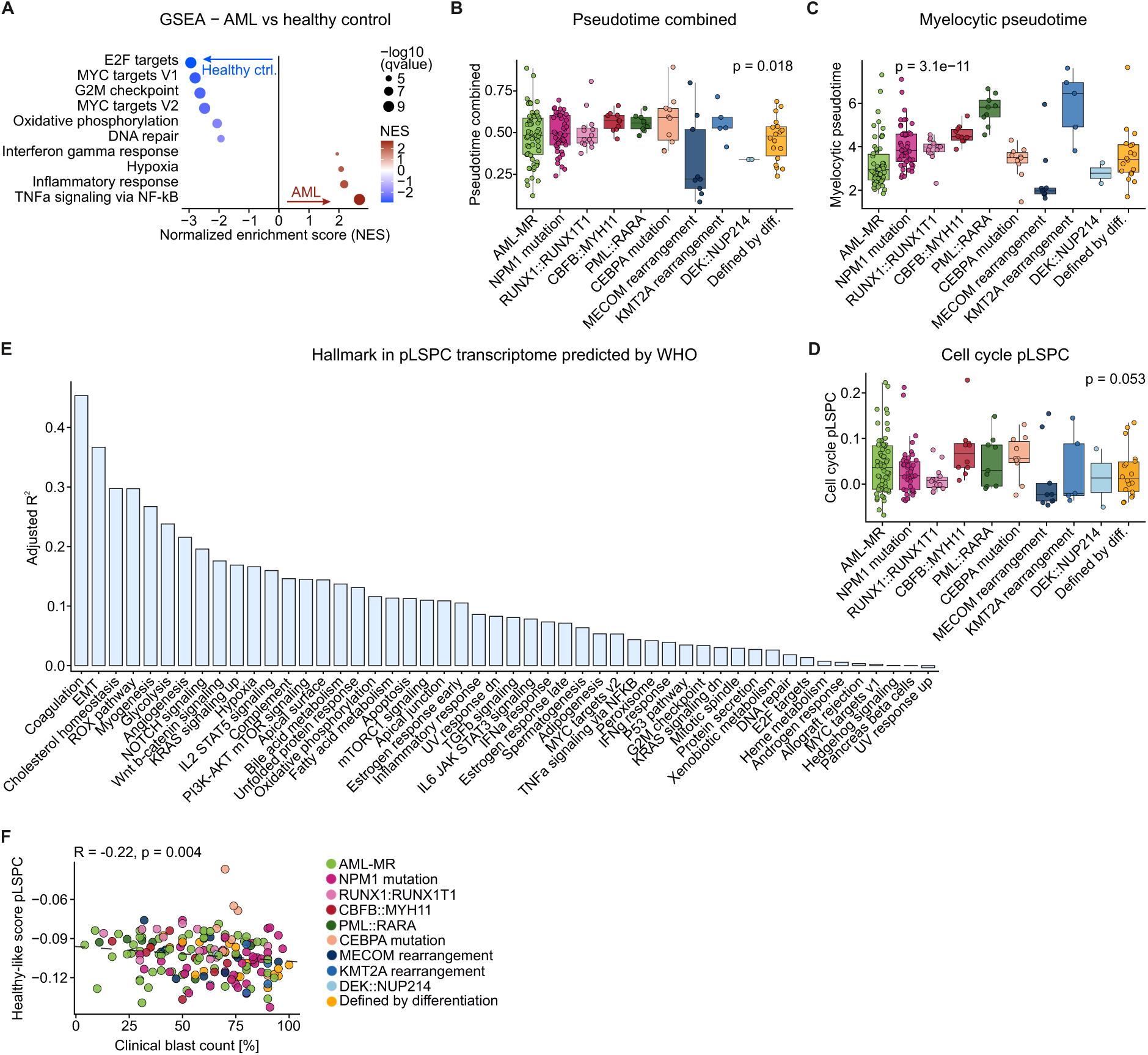
Genetic subtype-associated variation and the healthy-vs-AML transcriptional axis. **(A)** Gene set enrichment analysis (GSEA) of Hallmark pathways enriched in AML samples versus healthy controls (ctrl.). Pseudobulk expression was modeled with DESeq2, including CLR-transformed major lineage composition as covariates. Normalized enrichment score (NES) indicates enrichment direction and magnitude; point size reflects −log10 q-value. **(B–D)** Pseudotime and cell cycle features across WHO subtypes. Group differences were assessed using the Kruskal–Wallis test.**(B)** Combined pseudotime derived from percentile-based integration of lineage-specific pseudotimes projected from a healthy bone marrow reference. **(C)** Myelocytic pseudotime. **(D)** Cell cycle score in the pLSPC compartment (S-phase gene set). **(E)** Hallmark pathway scores in the pLSPC compartment predicted by WHO subtype. For each Hallmark gene set, linear regression was used to model pathway score as a function of WHO subtype. Bar height shows adjusted R², indicating the strength of association between WHO subtype and pathway activity. **(F)** Correlation between clinical blast count and healthy-like score. Spearman’s rank correlation coefficient is indicated. Each point represents an individual patient sample, colored by WHO 2022 classification. Dashed lines show ordinary least squares fit for visualization.

**Supplementary Figure 6.**
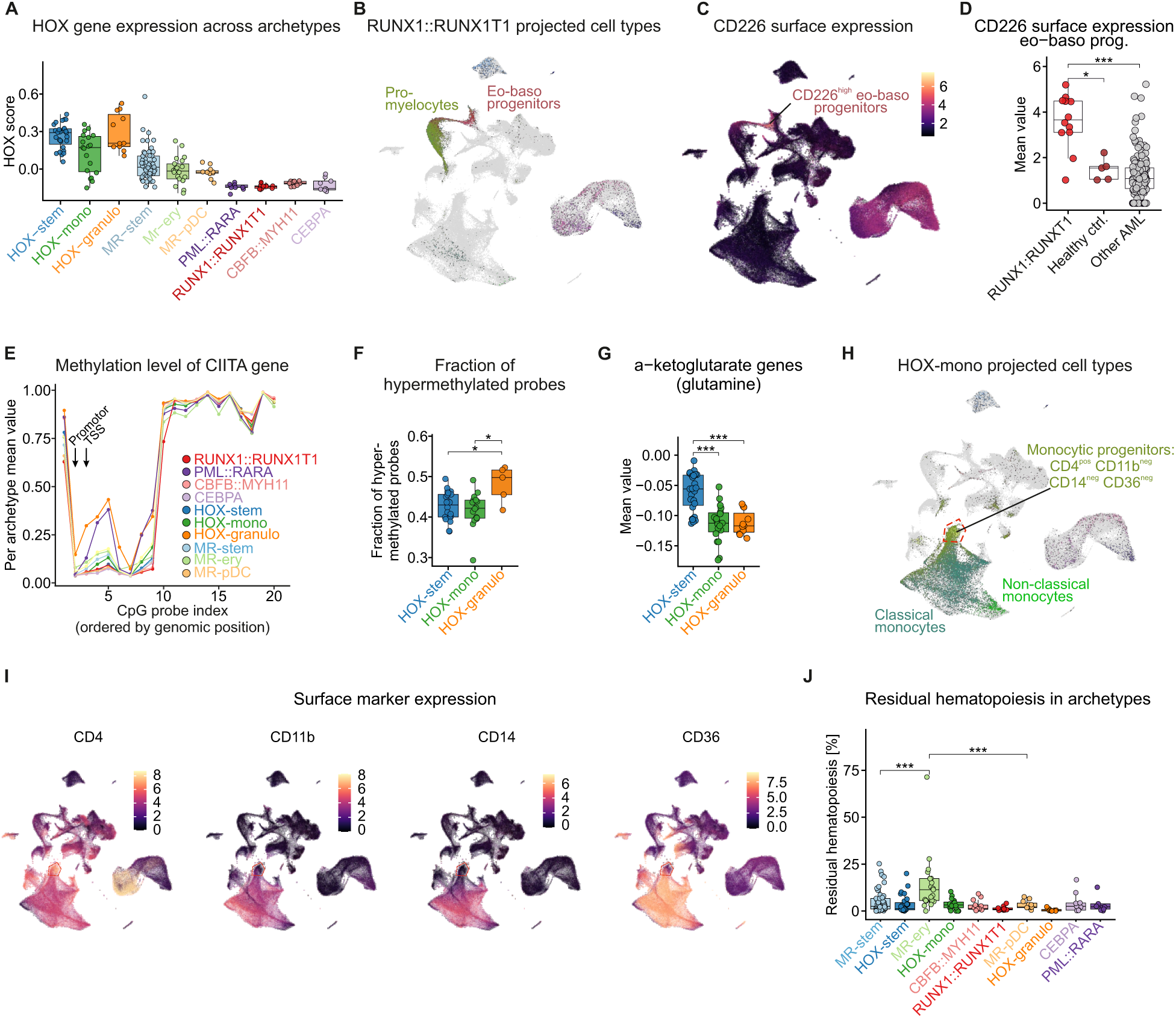
Molecular, epigenetic, and phenotypic features of AML archetypes. **(A)** HOX gene set expression across AML archetypes. Score was computed from canonical HOXA/B genes (HOXA3–10, HOXB3–6). **(B)** RUNX1::RUNX1T1 AML cells shown on the single-cell RNA-seq UMAP, colored by projected cell type. Promyelocytic and eosinophil– basophil progenitor populations are highlighted as the dominant lineages. **(C)** Single-cell surface expression of CD226 shown on the integrated UMAP. CD226^high^ eosinophil–basophil (eo-baso) progenitors are highlighted. **(D)** CD226 surface expression in eosinophil–basophil progenitors across indicated patient groups and healthy controls (ctrl.). **(E)** DNA methylation across the CIITA locus in TCGA AML samples. Archetype labels were assigned by estimating archetypes in patient-matched bulk RNA-seq data. Methylation probes are ordered by genomic position on the x-axis, and the y-axis shows the mean methylation level for each probe within each archetype. Promoter and transcription start site (TSS) regions are indicated. **(F)** Fraction of hypermethylated probes across HOX archetypes. For each sample, the fraction of assayed probes with β-value > 0.8 was calculated and compared across the indicated archetypes. Points represent individual samples. **(G)** Genes involved in α- ketoglutarate production and glutamine-driven anaplerosis were aggregated into a gene set score and quantified in the pLSPC compartment. Scores were compared across the indicated HOX archetypes. **(H)** HOX-mono archetype cells displayed on the single-cell RNA-seq UMAP and colored by projected cell type. Cells with a monocytic leukemic stem cell (mLSC) phenotype, defined as CD4^pos^CD11b^neg^CD14^neg^CD36^neg 5^, are highlighted. **(I)** Surface expression of the mLSC-defining markers (CD4, CD11b, CD14, and CD36). **(J)** Percentage of residual hematopoiesis across AML archetypes. Differences among the myelodysplasia-related (MR) archetypes are highlighted. **(D, F, G, J)** P values were determined using the Wilcoxon rank-sum test. Significance levels are indicated as follows: * p < 0.05, ** p < 0.01, *** p < 0.001 , **** p < 0.0001.

**Supplementary Figure 7.**
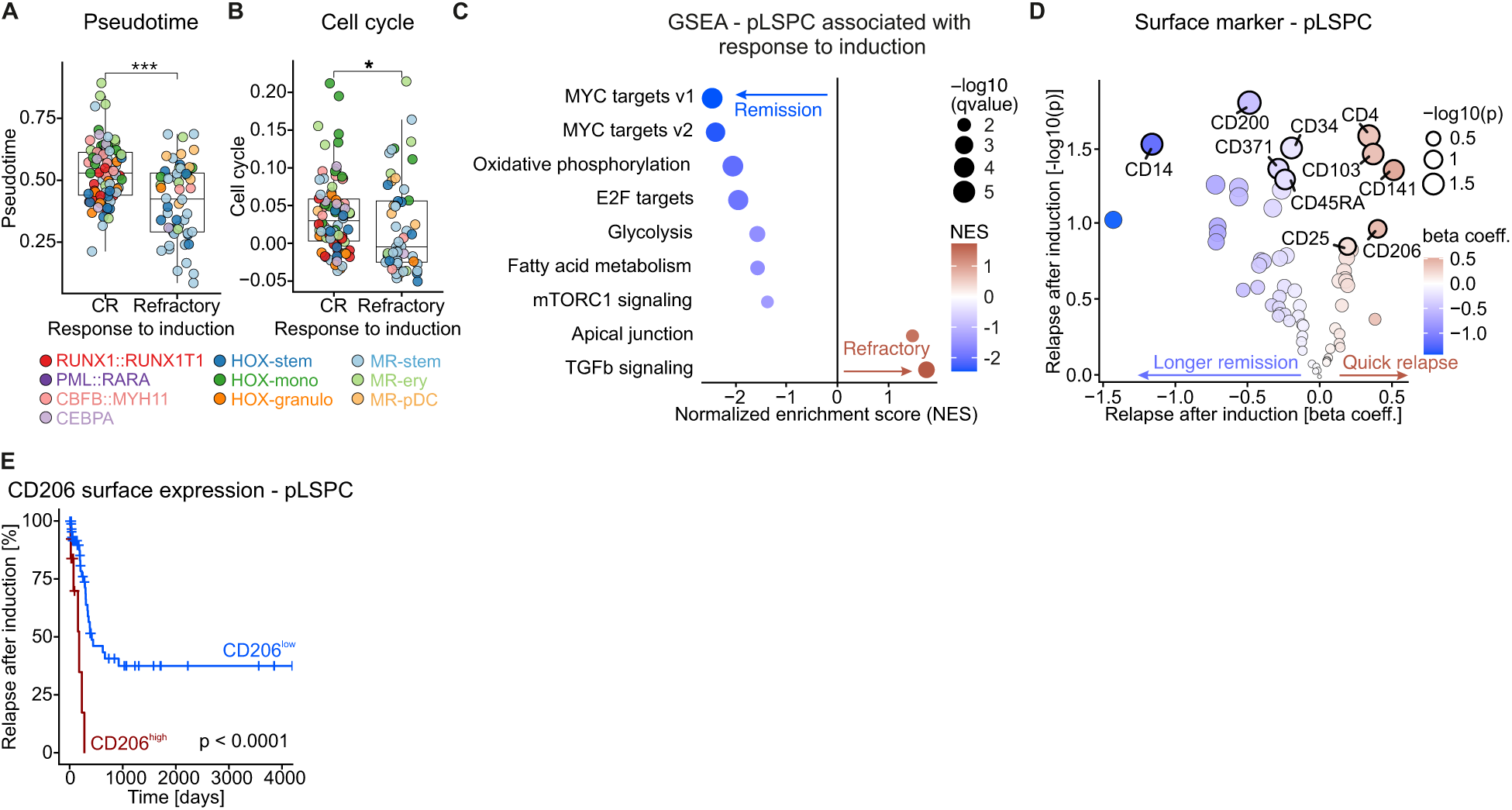
Features associated with response and relapse after induction therapy. (A,B) Association of pseudotime **(A)** and cell cycle activity **(B)** with response to induction therapy. Each point represents an individual sample and is colored by AML archetype. P values were calculated using the Wilcoxon rank-sum test. **(C)** Gene set enrichment analysis (GSEA) of pathways associated with response to induction using gene expression profiled in the pLSPC compartment (analogous to Figure 5I). The x-axis shows the normalized enrichment score (NES), with positive values indicating association with refractory disease and negative values indicating association with remission. Point size reflects statistical significance (−log10 q-value). **(D)** Volcano plot showing associations between surface marker expression in the pLSPC compartment and relapse after induction therapy. Surface markers were filtered to retain antibodies with Pearson correlation p < 0.05 between patient-level AbSeq abundance and matched RNA expression in pLSPCs, to reduce potential effects of nonspecific background or ambient signal. The x-axis shows the Cox regression beta coefficient and the y-axis shows the −log10 p-value. Point color indicates effect direction and magnitude, and point size indicates significance. The top five markers by p-value are labeled for each direction of association. **(E)** Relapse after induction therapy stratified by CD206 expression in the pLSPC compartment. High and low groups were defined using data-derived survival cutpoints selected to maximize separation of survival outcomes, and statistical significance was assessed using a log-rank test. Significance levels are indicated as follows: * p < 0.05, ** p < 0.01, *** p < 0.001 , **** p < 0.0001.

**Supplementary Figure 8.**
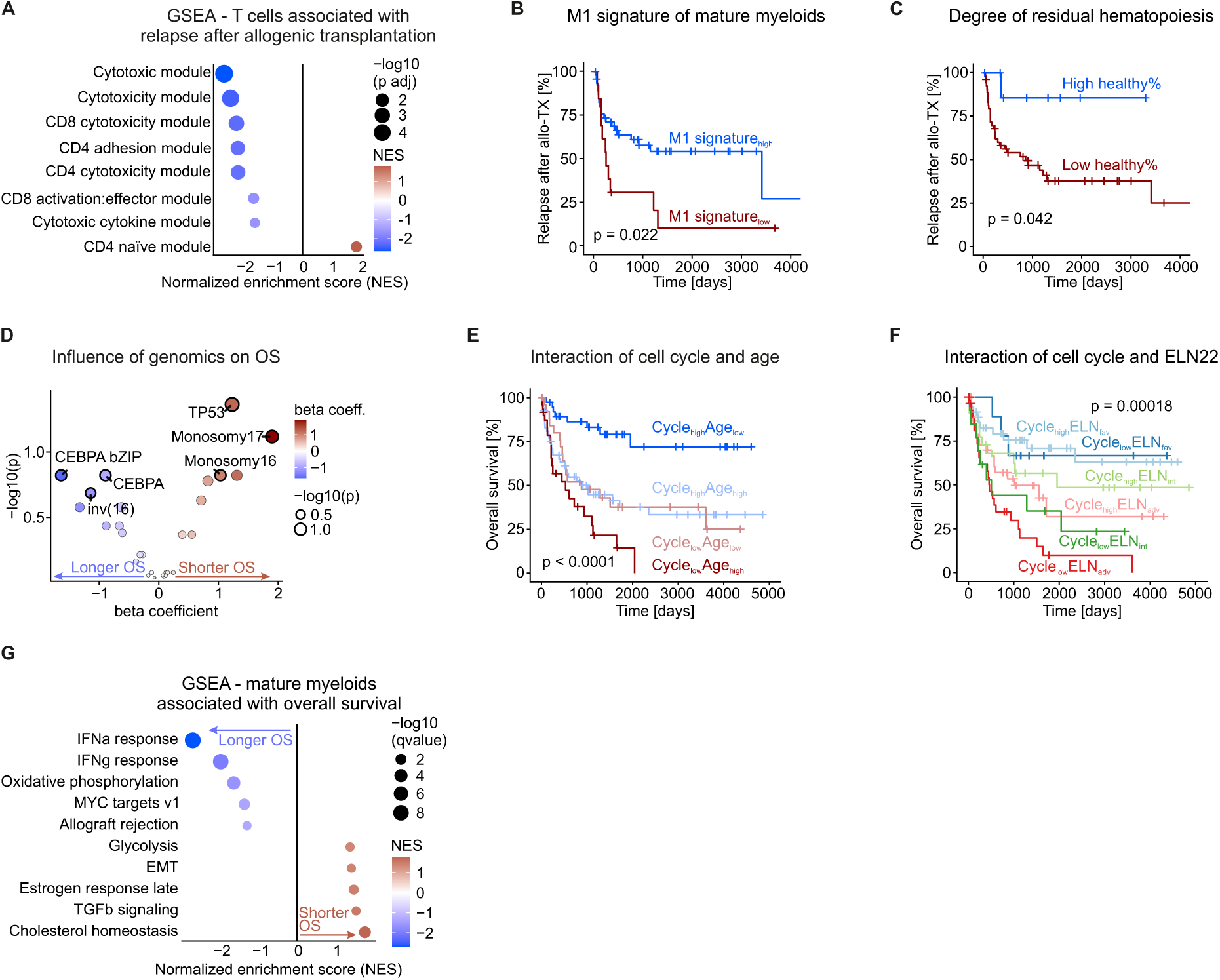
Determinants of post-transplant relapse and overall survival. **(A)** Patients were stratified into T cell–high and T cell–low groups using the data-derived cutpoint shown in Figure 6b. Lymphoid compartment pseudobulks were generated from T and NK/T cell populations, and differential expression between T cell–high and T cell–low patients was modeled with DESeq2. Genes were ranked by the DESeq2 Wald statistic and analyzed by GSEA using curated T cell gene sets. The x-axis shows the normalized enrichment score (NES), and point size reflects statistical significance (−log10 adjusted p-value). Gene sets not manually curated: cytotoxic module^6^, cytotoxicity module^7^, cytotoxic cytokine module^8^ **(B,C)** Relapse after allogeneic stem cell transplantation stratified by mature myeloid M1 signature score **(B)** and residual hematopoiesis **(C)**. **(D)** Genetic associations with overall survival. Cox proportional hazards models were used to test individual genomic features. The x-axis shows the Cox regression coefficient, with positive values indicating association with shorter overall survival and negative values indicating association with longer overall survival. The y-axis shows statistical significance as −log10 p-value. **(E,F)** Overall survival stratified by combined clinical and molecular features: cell cycle activity and age **(E)**, and cell cycle activity and ELN22 genetic risk **(F)**. **(G)** Gene set enrichment analyses (GSEA) of Hallmarks linked to overall survival in mature myeloid compartment (analogous to Figure 5I); normalized enrichment score (NES) indicates direction and strength of association, and point size reflects statistical significance (−log10 adjusted p-value). **(B, C, E, F)** High and low groups were defined using data-derived survival cutpoints selected to maximize separation of survival outcomes, and statistical significance was assessed using log-rank tests.

